# Zoledronic acid improves bone quality and muscle function in a high bone turnover state

**DOI:** 10.1101/2023.06.01.543305

**Authors:** Trupti Trivedi, Mohamed Manaa, Sutha John, Steven Reiken, Sreemala Murthy, Gabriel M. Pagnotti, Neha S. Dole, Yun She, Sukanya Suresh, Brian A. Hain, Jenna Regan, Rachel Ofer, Laura Wright, Alex Robling, Xu Cao, Tamara Alliston, Andrew R. Marks, David L. Waning, Khalid S. Mohammad, Theresa A. Guise

## Abstract

**SUMMARY:** Zoledronic acid (ZA) prevents muscle weakness in mice with bone metastases; however, its role in muscle weakness in non-tumor-associated metabolic bone diseases and as an effective treatment modality for the prevention of muscle weakness associated with bone disorders, is unknown. We demonstrate the role of ZA-treatment on bone and muscle using a mouse model of accelerated bone remodeling, which represents the clinical manifestation of non-tumor associated metabolic bone disease. ZA increased bone mass and strength and rescued osteocyte lacunocanalicular organization. Short-term ZA treatment increased muscle mass, whereas prolonged, preventive treatment improved muscle mass and function. In these mice, muscle fiber-type shifted from oxidative to glycolytic and ZA restored normal muscle fiber distribution. By blocking TGFβ release from bone, ZA improved muscle function, promoted myoblast differentiation and stabilized Ryanodine Receptor-1 calcium channel. These data demonstrate the beneficial effects of ZA in maintaining bone health and preserving muscle mass and function in a model of metabolic bone disease.

**Context and significance:** TGFβ is a bone regulatory molecule which is stored in bone matrix, released during bone remodeling, and must be maintained at an optimal level for the good health of the bone. Excess TGFβ causes several bone disorders and skeletal muscle weakness. Reducing excess TGFβ release from bone using zoledronic acid in mice not only improved bone volume and strength but also increased muscle mass, and muscle function. Progressive muscle weakness coexists with bone disorders, decreasing quality of life and increasing morbidity and mortality. Currently, there is a critical need for treatments improving muscle mass and function in patients with debilitating weakness. Zoledronic acid’s benefit extends beyond bone and could also be useful in treating muscle weakness associated with bone disorders.

## INTRODUCTION

Bone metabolism is a highly dynamic process that involves interactions between bone cells (Hadjidakis and Androulakis, 2006, Siddiqui and Partridge, 2016) as well as between bone and skeletal muscle (Brotto and Bonewald, 2015, Regan et al., 2016, Karsenty and Mera, 2018, Regan et al., 2017b). Bone-muscle cross-talk occurs via mechanical forces that influence these tissues and endocrine signaling, and pathological bone loss can negatively impact muscle function (Regan et al., 2017b, Regan et al., 2017a, Waning et al., 2015, Waning and Guise, 2014). Manifestations of bone loss and muscle weakness are present under conditions such as aging (Edwards et al., 2015, Wagner et al., 2018), osteoporosis (Pham et al., 2017), sex steroid deficiency (Sipila et al., 2013, Wright et al., 2017, Storer et al., 2012), Paget disease (Hayashi, 2013), and cancer treatment (Storer et al., 2012, Wright et al., 2017, Hain et al., 2019). Simultaneous bone loss and muscle weakness are associated with limited mobility, falls, fractures, and disability (Pham et al., 2017, Ferrucci et al., 2014, Tarantino et al., 2015, Scott et al., 2017).

Zoledronic acid (ZA) is a U.S. Food and Drug Administration-approved bone antiresorptive agent for treating primary and secondary osteoporosis, cancer-associated bone diseases, and Paget disease (Rakel et al., 2011, Guise, 2006, Brown and Guise, 2007, Coleman et al., 2010, Corral-Gudino et al., 2017, Coleman et al., 2011). ZA inhibits osteoclast proliferation and induces osteoclast apoptosis, preventing bone resorption (Rogers et al., 2000). In addition, ZA promotes osteoblast differentiation and increases bone mineralization (Reinholz et al., 2000). Despite the use of ZA to treat various bone diseases for decades, its effects on muscle function in patients with nonmalignant bone diseases are not established.

Transforming growth factor β (TGFβ) is synthesized as a large precursor molecule by osteoblasts and embedded in the mineralized bone matrix but remains in an inactive or latent state (Yin et al., 1999). Active TGFβ is released into the circulation during osteoclast-mediated bone resorption and regulates bone remodeling via pleiotropic effects on osteoclasts and osteoblasts and their precursors as well as osteocytes (Khalil, 1999). Previously, using seven different mouse models of bone metastases and a model of multiple myeloma in bone, we showed that osteoclast-mediated bone loss resulted in the systemic release of bone-derived TGFβ, causing skeletal muscle weakness, which was prevented by treatment with ZA (Waning et al., 2015). ZA also has antitumor effects (Morgan and Lipton, 2010, Clezardin, 2002), raising the possibility that prevention of muscle weakness in models of bone metastases by ZA could be the result of blocking the release of bone-derived TGFβ and reducing the tumor burden. However, muscle weakness with bone metastases may be a mixed effect of bone-derived TGFβ (Waning et al., 2015) and bone-independent effects of a tumor on muscle (Hogan et al., 2018, Zhang et al., 2017). Besides its prevention of bone resorption, ZA has been reported to have antitumor effects (Morgan and Lipton, 2010, Clezardin, 2002) which raising the possibility that prevention of muscle weakness in models of bone metastases by ZA could be the result of blocking release of bone-derived TGFβ and reduction of tumor burden.

Metabolic bone diseases not associated with cancer may yield important understanding of the specific TGFβ-mediated effects on skeletal muscle weakness. Camurati-Engelmann disease (CED), or progressive diaphyseal dysplasia, is an autosomal dominant disorder caused by point mutations in the latent peptide region of TGFβ, leading to pathological bone remodeling. CED is characterized by severe skeletal abnormalities due to extreme bone turnover and increased TGFβ signaling. A mouse model of CED (Tang et al., 2009) mimics these clinical features, making it suitable for studying the sssystemic effects of excess bone-derived TGFβ in the absence of bone metastases. Given the simultaneous occurrence of bone loss and muscle weakness in CED mice, we examined the muscle weakness and evaluated the potential of ZA for treating the severe bone defects and simultaneously improving muscle function. In addition to improvement of bone volume and muscle mass, improved musculoskeletal function is essential as a therapeutic strategy for both tissues (Schuit et al., 2004, Sornay-Rendu et al., 2007, Kim and Kang, 2017, Amthor et al., 2007, Wagner et al., 2018, Loenneke et al., 2019). Mobility limitations due to bone destruction and its effects on muscle occur in almost all patients suffering from bone diseases. Recognizing and preventing the muscle weakness that occurs as a consequence of increased bone turnover will reduce morbidity and mortality.

The treatment strategy with ZA for bone disorders is well established. However, ZA treatment duration, treatment modalities (preventative or therapeutic) in bone disorders associated with muscle weakness is unknown. It is also unknown if ZA has any direct effects on the regulation of skeletal muscle physiology. To identify the extra-skeletal effects of ZA in muscle we hypothesized that preventative ZA treatment increase muscle mass and function in mice with extreme bone remodeling and excess TGFβ. In the present prospective study, we showed that bone-derived TGFβ is a key factor causing muscle weakness from high bone turnover in the absence of cancer. Excess TGFβ reduces the skeletal muscle myofiber cross-sectional area, causes a muscle fiber type shift from oxidative to glycolytic, and disrupts the osteocyte lacunocanalicular network. Treatment with ZA improves each of these effects of excess TGFβ and results in increased bone strength and improved muscle function. Our studies have highlighted the importance of improved overall musculoskeletal function with ZA treatment in mice with CED.

## RESULTS

### Bone loss causes muscle weakness in mice with CED

To elucidate the role of bone-derived TGFβ in skeletal muscle in a nonmalignant model of altered bone metabolism, we used a mouse model of CED because it recapitulates many of the clinical features of accelerated bone remodeling of human bone diseases. CED mouse model was generated by osteoblast-specific expression of CED mutant TGF-β1 exclusively in bone. To determine whether the increased bone remodeling in CED is associated with skeletal muscle weakness, we measured muscle contractility by determining the specific force-frequency curve of the extensor digitorum longus (EDL) muscle in female C57BL/6 mice with CED and their wild-type (WT) littermates. At 6 weeks of age, we saw no difference in the EDL muscle contractility, but by 8 weeks and continuing at 12 weeks of age, mice with CED exhibited lower EDL muscle contractility than did the WT mice (Figure 1A). We previously reported that EDL muscle contractility was lower in male mice with CED than in their WT littermates (Waning et al., 2015). Because we observed muscle weakness in mice with CED at 8-12 weeks of age, we selected mice in this age group for further experiments.

**Figure 1.**
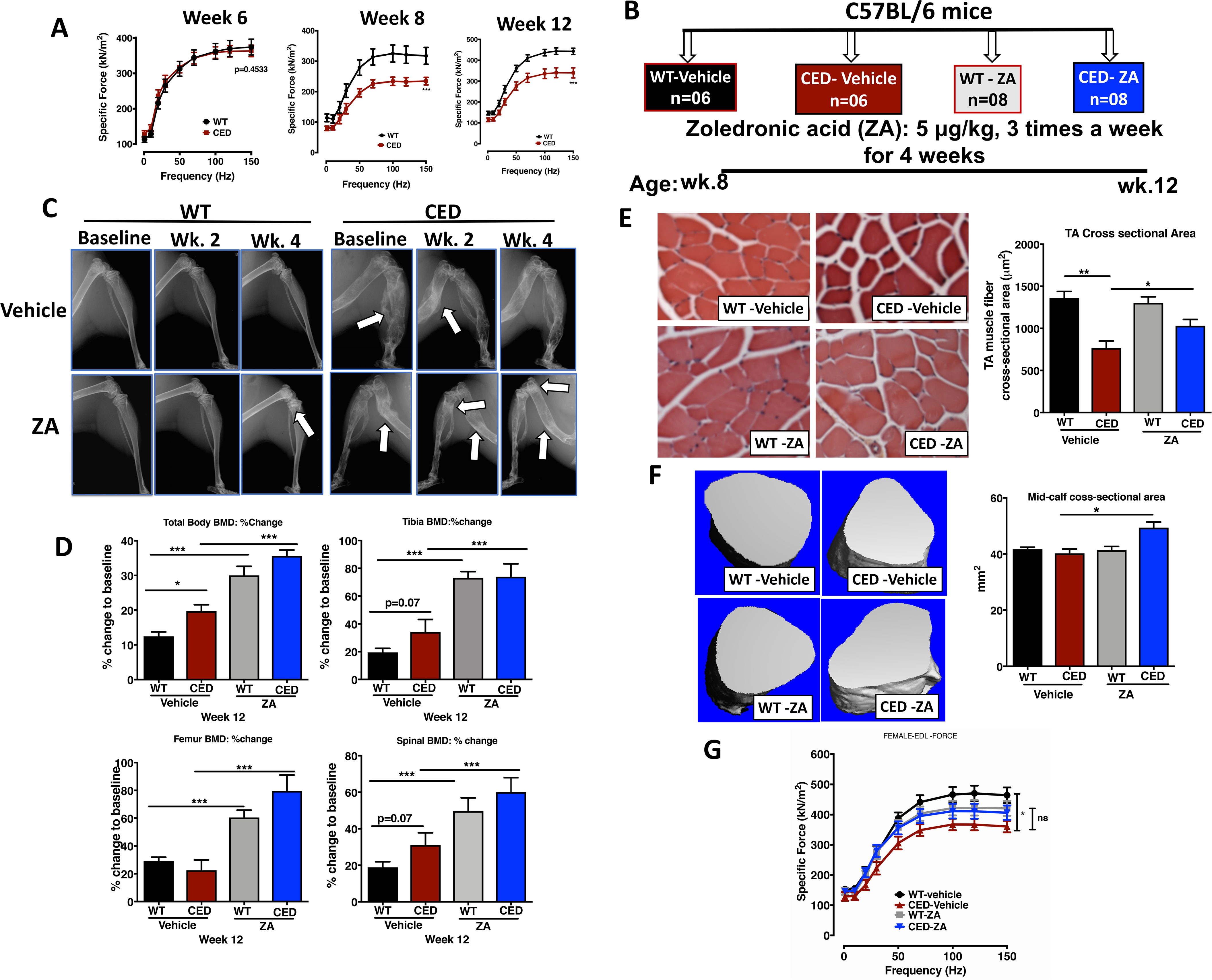
Assessment of bone changes and muscle weakness in WT mice and mice with CED in response to 4-week ZA treatment. (A) *Ex vivo* EDL muscle contractility in WT mice and mice with CED at the age of 6, 8, and 12 weeks and normalized according to muscle size. (B) The experimental design, showing the groups of mice and ZA administration strategy. (C) Digital x-rays (KUBTEC XPERT 80) of ZA treatment effects in the two mouse groups monitored at baseline (before treatment) and after 2 and 4 weeks of treatment. In the upper panels, the arrows indicate diaphyseal dysplasia in mice with CED. In the lower panels, the arrows indicate progressive increase in bone density at metaphysis in WT mice and mice with CED. The arrow at diaphysis in ZA-treated mice with CED indicates changes in bones with the combination of ZA and/or abnormal bone remodeling. (D) BMD in 12-week-old mice as determined using DXA after 4 weeks of treatment. Differences in BMD are expressed as percentage change from baseline. Whole-body BMD and the BMD at the proximal tibiae, distal femora, and spine were measured. (E) TA muscle fiber diameters determined using hematoxylin and eosin staining of muscle sections and evaluated to measure muscle cross-section areas in response to ZA treatment. (F) *In vivo* micro-CT assessment of muscle mid-calf cross-sectional areas following 4-week ZA treatment in WT mice and mice with CED. (G) *Ex vivo* EDL muscle contractility assessment in WT mice and mice with CED after 4 weeks of ZA treatment. EDL force was normalized according to muscle size. Data are expressed as mean (± SEM) differences in muscle function was analyzed using two-way analysis of variance (ANOVA) with Bonferroni’s multiple comparison testing. *p < 0.05, **p<0.01, and ***p<0.001. Differences in all other parameters were determined using one-way ANOVA with Tukey’s multiple comparison test or an unpaired t-test. *p < 0.05, **p<0.01, and ***p<0.001. Differences between vehicle-treated WT mice and mice with CED and between vehicle- and ZA-treated mice with CED are presented.

### ZA treatment for 4 weeks increases bone mineral density and muscle mass but not muscle function in mice with CED

We next investigated whether inhibition of bone resorption by ZA treatment for 4 weeks also improves muscle function in mice with CED. To determine whether ZA alone has any effect on muscle in healthy mice, we gave WT mice two doses of ZA (2.5 μg/kg once a week or 5 μg/kg three times a week) for 4 weeks. Both doses increased trabecular bone volume (Figures S1A and S1B) without affecting muscle function or mass (Figures S1C and S1D), demonstrating that ZA does not affect skeletal muscle in the absence of abnormal bone metabolism. To account for the extremely high levels of bone turnover in mice with CED, in the next experiment, we treated WT mice and those with CED 5 μg/kg ZA three times a week for 4 weeks (Figure 1B). Due to increased bone remodeling, mice with CED had overall higher total bone mineral density (BMD) than did WT mice. Also, ZA produced higher BMD in the tibia, femur, and lumbar spine in both groups of mice than in corresponding vehicle-treated control groups (Figures 1C and 1D). WT mice had two-fold higher tibialis anterior (TA) myofiber diameters than did mice with CED, and ZA significantly increased TA myofiber diameters in mice with CED but not WT mice (Figure 1E). Consistent with the increase in myofiber diameter, ZA also increased the mid-calf cross sectional area in CED mice (Figure 1F). In mice with CED, 4 weeks of ZA treatment resulted in an increase in EDL muscle contractility, but it did not reach statistical significance (Figure 1G), suggesting that a longer period of ZA treatment might improve muscle function.

### ZA treatment for 10 weeks increases BMD in mice with CED

Despite the improvements in BMD in mice with CED after 4 weeks of treatment with ZA, an extended treatment time may be required to sufficiently block TGFβ release from bone to an extent that can significantly improve skeletal muscle function. To examine whether prolonged ZA treatment can substantially increase muscle function in mice with CED, we administered ZA to them for 10 weeks. We started treatment when the mice were 7 weeks of age to ensure treatment of bone loss before the onset of muscle weakness. Our results demonstrated that after 10 weeks of treatment, ZA reduced serum TGFβ concentrations in mice with CED (Figure 2A) but did not change TGFβ concentrations in WT mice. In response to ZA exposure, we observed a time-dependent increase in radiographic bone density in both groups of mice (Figure 2B). ZA treatment improved the severity of the diaphyseal dysplasia in CED mice but was not completely normalized, with less severe lesions observed on x-ray. This is likely due to continuous release of active bone-derived TGFβ in these mice, which stimulates bone turnover. Furthermore, we assessed BMD at the proximal tibia and distal femur (a site of normal active bone remodeling) and the entire tibia and femur (to include a site of dysplasia and abnormal bone remodeling visualized in x-rays of mice with CED). After 10 weeks, total body BMD was higher in mice with CED than in WT mice after treatment with ZA but similar in the two groups after treatment with a vehicle. ZA-based treatment produced higher BMD in the proximal tibia, distal femur, whole tibia, whole femur, and lumbar spine than did the vehicle in both WT mice and mice with CED. However, BMD in the tibia, femur, and spine did not differ between the WT and CED groups after treatment with the vehicle. Only at the distal femur did we see a trend of higher BMD in mice with CED than in WT mice given the vehicle (Figure 2C).

**Figure 2.**
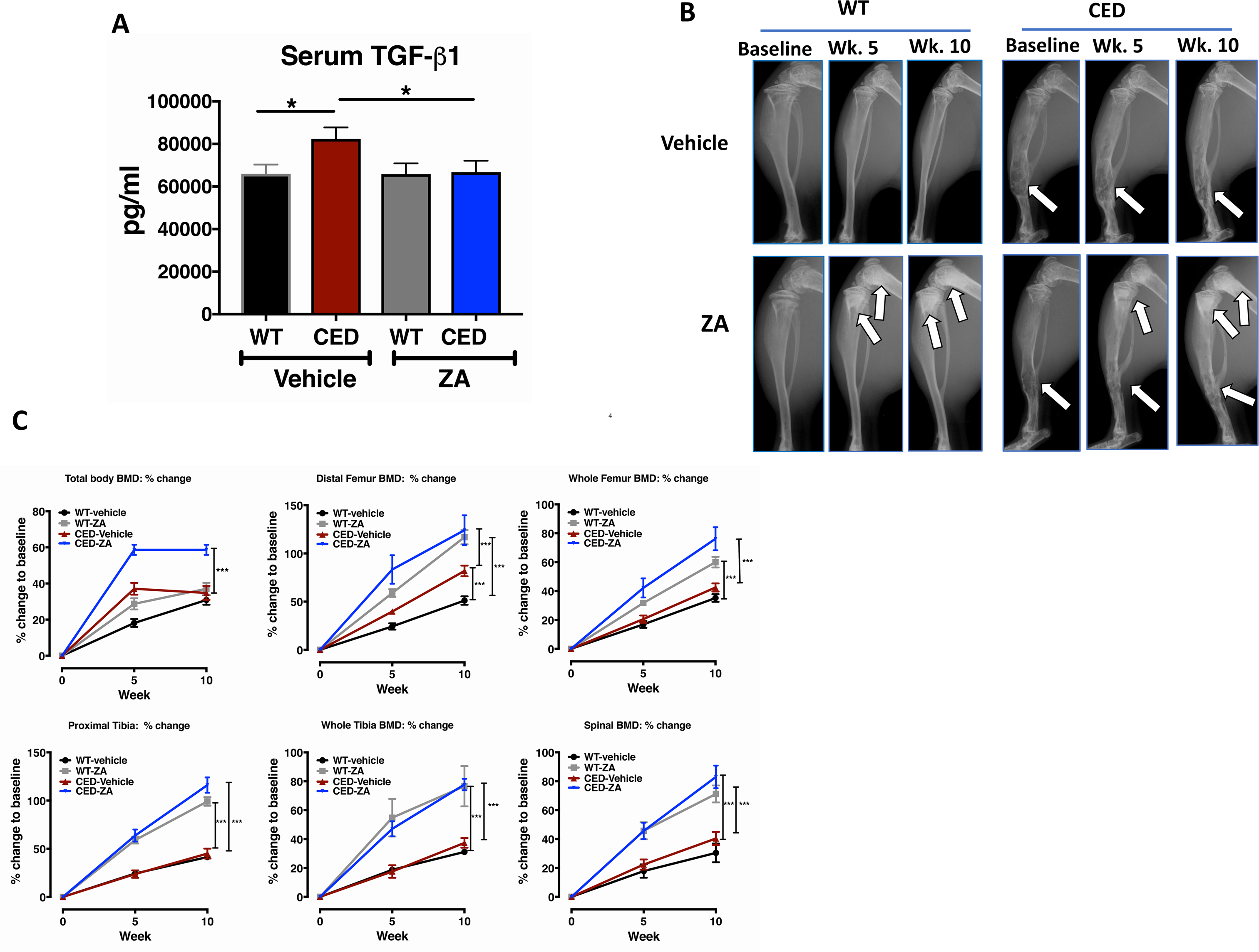
Ten-week ZA treatment effects in WT mice and mice with CED determined using x-rays and DXA. (A) ZA-based treatment was started in 7-week-old WT C57BL/6 mice and C57BL/6 mice with CED to start blocking bone resorption before muscle weakness began. The mice were placed in four groups and given a vehicle (sterile phosphate-buffered saline [PBS]) or ZA for 10 weeks. Serum was collected from the mice after 10 weeks of treatment and assayed for TGFβ1 using enzyme-linked immunosorbent assay (ELISA). (B) ZA treatment effects were monitored from baseline (prior to treatment) to weeks 5 and 10 (before euthanasia) using radiography. The arrows indicate progressive increases in bone density at metaphysis in WT mice in response to treatment with ZA. Ten-week treatment effects of ZA in CED mice were evident at metaphysis and diaphysis (mixed effects of abnormal bone remodeling and ZA) as indicated by arrows. (C) BMD assessed using DXA at baseline, week 5, and week 10 in whole-body scans (total body BMD) and at the vertebral column (L4-L6), proximal tibiae, whole tibiae, distal femora, and whole femora. Data are expressed as means (± SEM), and differences groups in DXA analysis were examined using two-way ANOVA with Bonferroni’s multiple comparison testing. *p < 0.05, **p<0.01, and ***p<0.001 Differences between vehicle-treated WT mice and mice with CED and between vehicle- and ZA-treated mice with CED are presented.

### ZA treatment improves the trabecular and cortical bone microarchitecture in mice with CED

Trabecular bone parameters were determined in the proximal tibia and L5 vertebra. CED vehicle-treated mice showed significantly reduced trabecular bone volume fraction (BV/TV %), trabecular number (Tb.N), connectivity density (Conn.D), and increased trabecular separation (Tb.Sp) and structure model index (SMI) compared to WT-vehicle treated mice. 10-week of ZA treatment significantly increased BV/TV %, Tb.N, Conn.D, and decreased Tb.Sp and SMI in CED mice both at tibia and L5. Trabecular thickness in CED vehicle-treated mice did not change in tibia but was reduced in L5 compared to WT-vehicle and improved in CED mice in response to ZA treatment (Figures 3A-D). Cortical porosity analysis was performed in the tibia 15 mm distally from midshaft. Cortical porosity (Ct.Po) and medullary area (Ma.Ar) were increased 8 fold in CED mice, but were largely corrected in response to ZA treatment compared to WT mice. Cortical bone area (Ct.Ar) and cortical thickness (Ct.Th) were reduced in CED mice compared to WT mice treated with vehicle. ZA treatment significantly increased Ct.Ar and Ct.Th in CED mice compared to untreated mice (Figures 3E-G).

**Figure 3.**
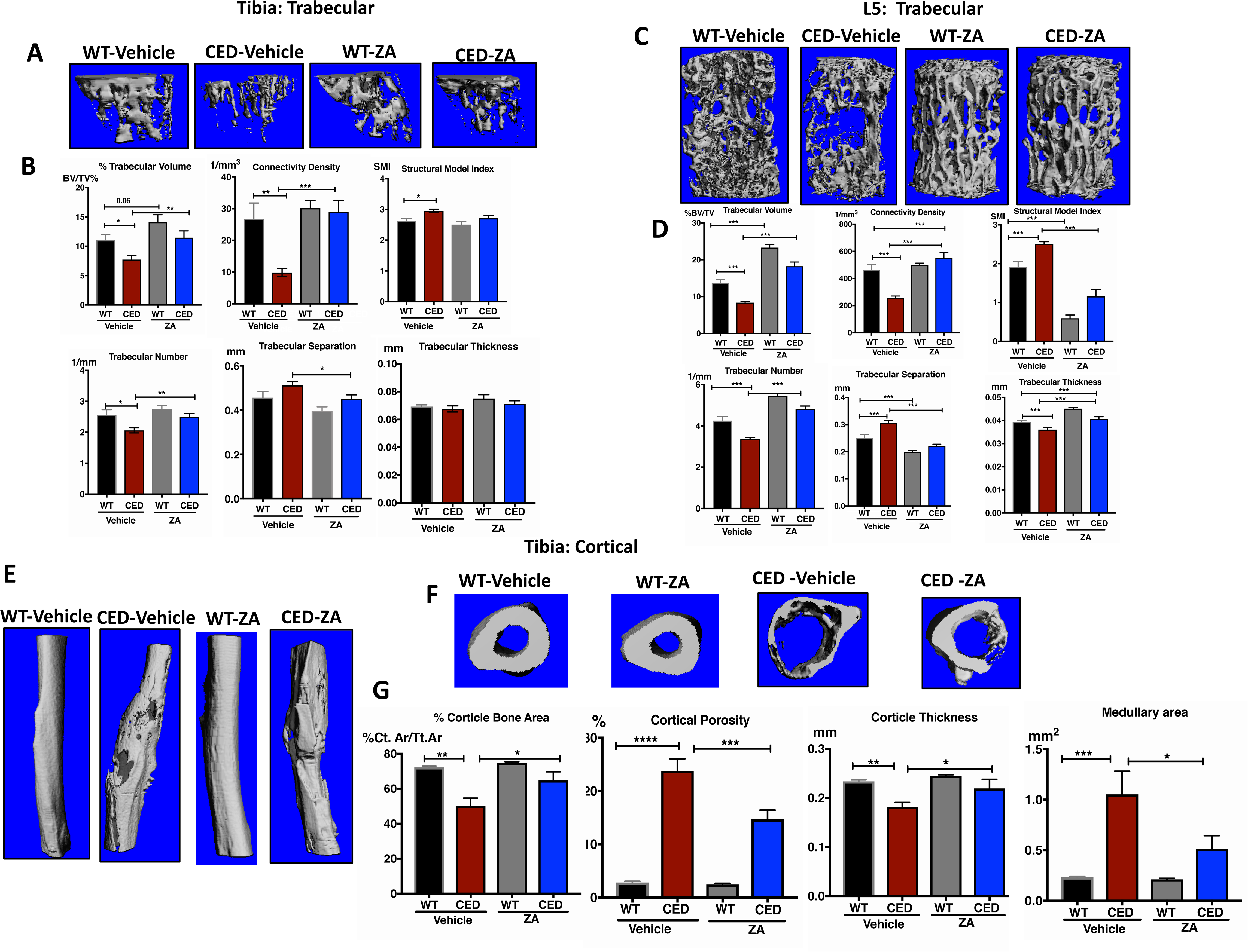
Bone microarchitecture assessed using micro-CT of *ex vivo* bone scans. (A) Trabecular bone microarchitecture in the proximal tibiae. Representative micro-CT reconstructions of the proximal tibiae are presented. (B) Differences in percent bone volume to tissue volume, connectivity density, structure model index, trabecular number, trabecular separation, and trabecular thickness. (C) Trabecular bone parameters measured in the L5 vertebral body as shown with micro-CT reconstructions of L5. (D) Trabecular parameters measured in L5 and differences between groups. (E) Representative images of cortical porosity of tibiae measured 15 mm distally from the midshaft by excluding the marrow space. (F) Separate analyses of cortical bone parameters were performed in the similar region as cortical porosity by including the marrow space. Representative images of cortical cross-sections of tibiae are shown. (G) Differences in cortical porosity (excluding the marrow space), cortical bone area, cortical thickness, and medullary area (including the marrow space). Data are expressed as means (± SEM), and differences between groups were determined using one-way ANOVA with Tukey’s multiple comparison test or an unpaired t-test. *p < 0.05, **p<0.01, ***p<0.001, Differences between vehicle-treated WT mice and mice with CED and between vehicle- and ZA-treated mice with CED are presented.

### Ten weeks of ZA treatment reduced the number of osteoclasts and overall rate of bone remodeling in mice with CED

Quantitative histomorphometry analysis revealed no difference in the number of tartrate-resistant acid phosphatase (TRAP)-positive osteoclasts at the bone surface of WT mice in response to ZA treatment. However, CED-vehicle treated mice show significantly increased number of TRAP positive osteoclasts at the bone surface compared to WT-vehicle treated mice. ZA treatment reduced TRAP-positive osteoclasts at the bone surface (Figure 4A). Consistent with previously reported effects of TGFβ on osteoblasts (Bonewald and Mundy, 1990), CED mice had a significantly increased number of osteoblasts at the bone surface, increased osteoid volume (OV), and osteoid surface (OS) in CED mice compared to WT mice with vehicle treatment. ZA treatment showed trend to reduce osteoblast numbers, OS and OV in both WT and CED mice (Figures 4B, C). Dynamic histomorphometry analysis was performed to determine the bone formation rate (BFR) over a seven-day interval. CED mice showed high bone formation rate compared to WT in vehicle treated groups. ZA treatment significantly reduced BFR, mineral apposition rate and mineralizing surface in CED mice. dynamic histomorphometry, TRAP and osteoid analysis collectively showed that ZA treatment in CED mice showed reduced bone formation rate, and reduced osteoclast and osteoblast numbers at the bone surface, indicating a reduction in the overall rate of bone remodeling (Figures 4B, C).

**Figure 4.**
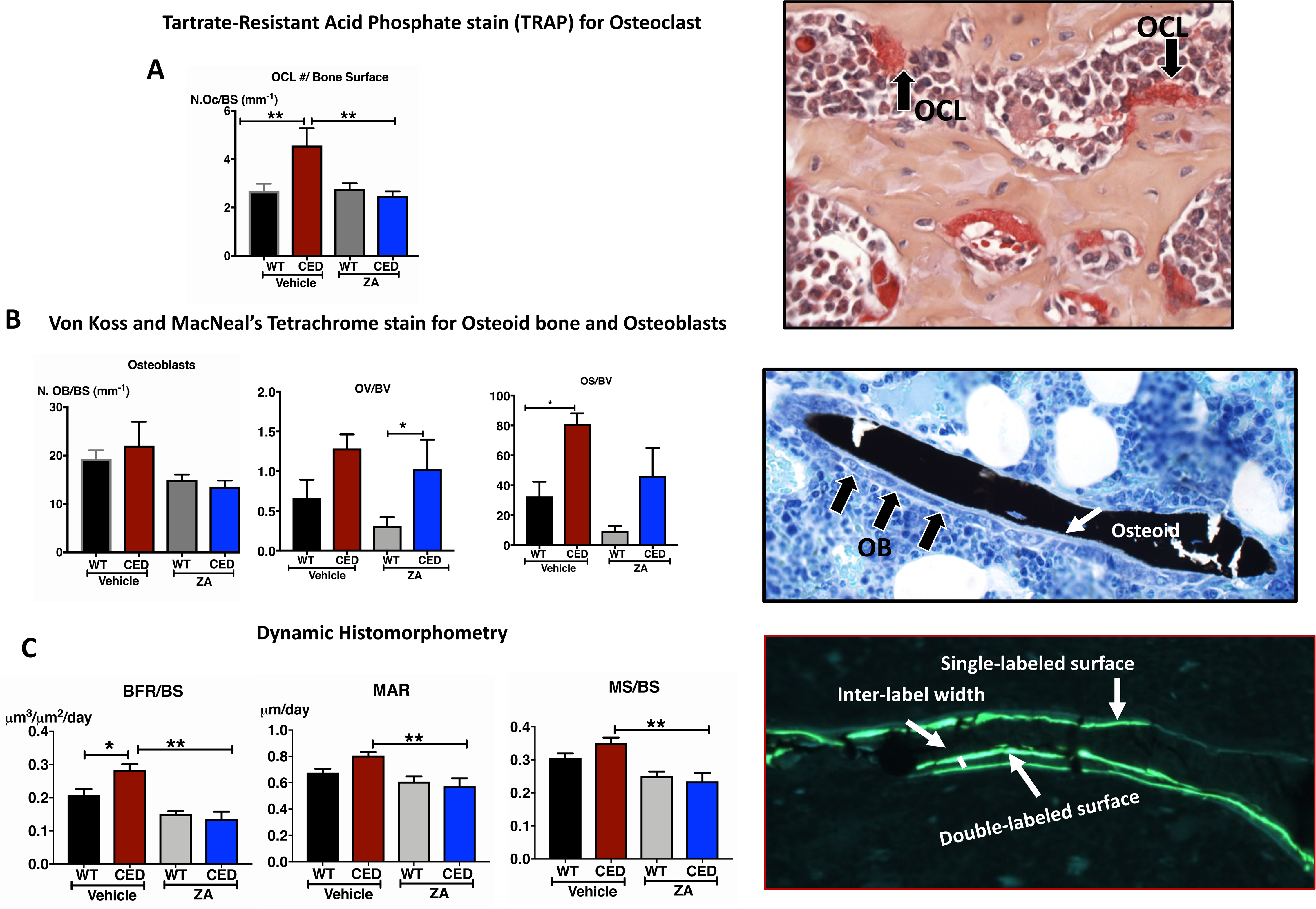
Histological assessment of bones. (A) TRAP-positive multinucleated osteoclasts in midsagittal sections of the tibiae quantitated at the bone surface (BS) at 20× magnification and expressed as the number of osteoclasts per millimeter of bone surface (OcN./mm). The black arrows show osteoblasts. (B) von Kossa/McNeal-stained undecalcified sections of distal femora. The white arrow indicates osteoid bone, and the black stain indicates mineralized bone. Analysis was performed at 20× magnifications to measure osteoid volume to bone volume, osteoid surface to bone surface, and osteoid width. (C) Dynamic histomorphometry of calcein-labeled femora was performed to assess the rate of bone formation. Calcein was injected 3 and 7 days prior to bone collection from mice. Calcein binds to newly formed bone to provide fluorescence. Dynamic histomorphometry was assessed at 20× magnification to determine the bone formation rate at the bone surface (BFR/BS), mineral apposition rate (MAR), and mineralized surface to bone surface (MS/BS). Data are expressed as means (± SEM), and differences were determined using one-way ANOVA with Tukey’s multiple comparison test or an unpaired *t*-test. *p < 0.05 and **p<0.01. Differences between vehicle-treated WT mice and mice with CED and between vehicle- and ZA-treated mice with CED are presented.

In CED mice, the overall increase in bone remodeling was reduced by ZA treatment. Despite this effect there was a significant increase in bone volume in CED-vehicle treated mice as seen by micro-CT possibly due to dominant bone formation effect of constitutively active TGFβ signaling in CED.

### ZA increases bone strength in mice with CED

To determine whether ZA-treatment improves bone biomechanical properties in mice with CED, we performed three-point bending tests on the femoral diaphyses and axial compression tests on the lumbar vertebral bodies. Similar to changes in bone microarchitecture determined by micro CT, with structure distortion indicated by arrows in representative femur in CED vehicle treated mice (Figure 5A), 3-point bending of the femur indicated reduced ultimate force, yield force, energy-to-yield and stiffness in CED mice compared to WT vehicle-treated mice (Figure 5B). Both CED and WT mice showed increased ultimate force, yield force, energy-to-yield and stiffness in response to ZA treatment and improved structure of the femur in CED mice (Figure 5A). L5 vertebral compression tests indicated lower ultimate force, yield force, energy to yield, and stiffness in vehicle-treated CED mice compared to vehicle-treated WT mice. In CED mice, ZA treatment increased the mechanical properties of L5 vertebrae, as indicated by increased ultimate force, yield force, energy to yield and stiffness, compared to vehicle-treated mice. (Figures 5C). In addition to improving bone microarchitecture, ZA treatment also increased mechanical properties in femora and L5 vertebrae (Figures 5B-C). Collectively, these data showed that ZA significantly increased bone volume and strength in mice with CED.

**Figure 5.**
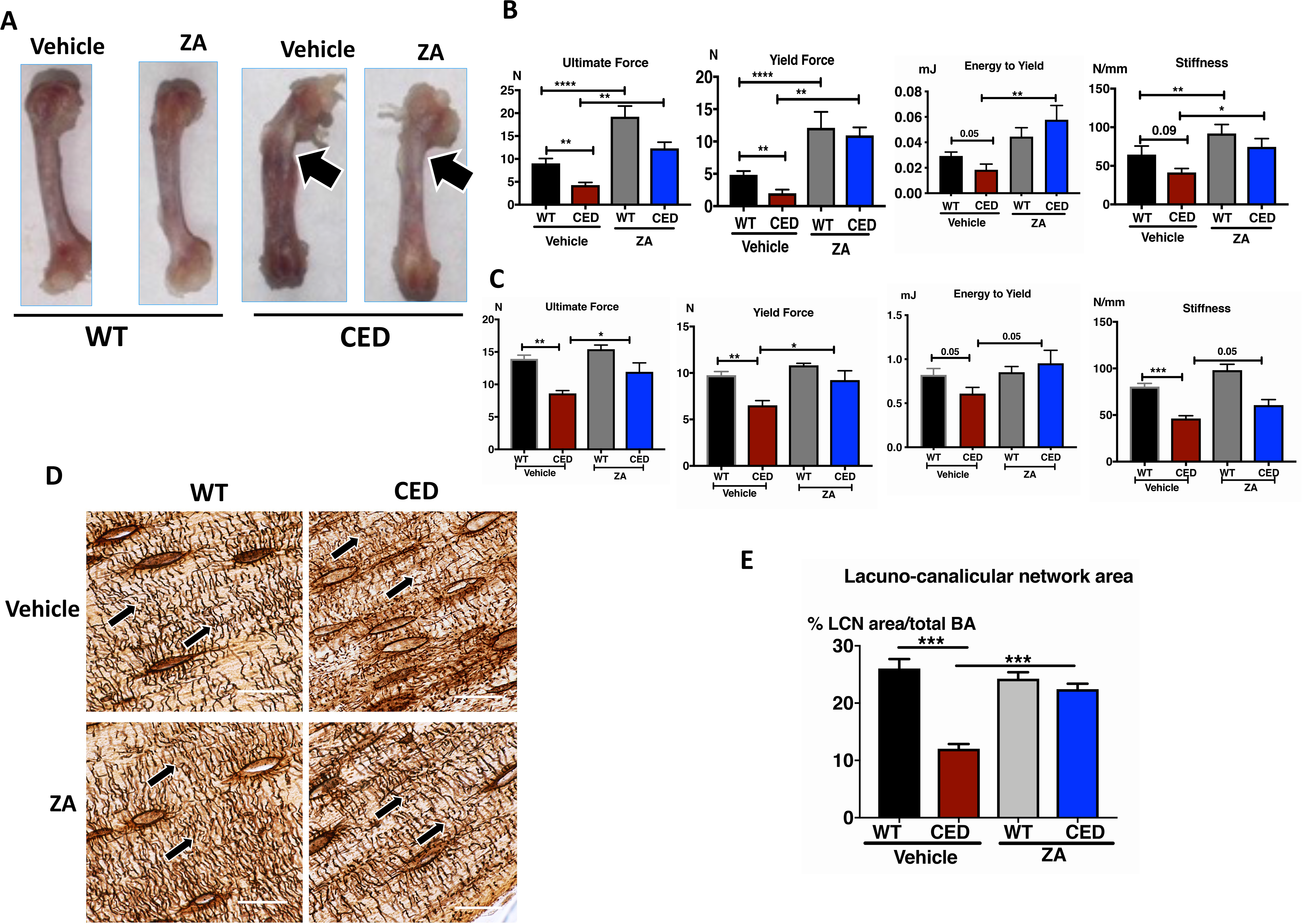
Osteocyte lacunocanalicular network analysis and bone mechanical properties Bone strength measured using three-point bending of the femur and compression testing of the L5 vertebra. (A) Representative images of femora from WT mice and mice with CED with and without ZA based treatment used for three-point bending. The arrows indicate the callus in a vehicle-treated mouse with CED and the absence of callus in the femur of a ZA-treated mouse with CED. (B) Differences in ultimate force, yield force, strain to yield, and stiffness were used to determine resistance to fracture. (C) Results of compression testing of the L5 vertebral body. Ultimate force, yield force, energy to yield, and stiffness were used to determine the strength of the bone based on force needed to break the vertebral body. Changes in L5 vertebral body strength are presented. (D) Representative osteocyte lacunocanalicular network images of cortical bone in paraffin embedded tibiae with Ploton silver staining. Degeneration of the LCN occurs when perilacuno/canalicular remodeling is suppressed, which compromises the mechanical integrity of the bone. The arrows indicate LCN areas. (E) Differences in the percent LCN area to total cortical bone area. Data are expressed as means (± SEM), and differences between groups were determined using one-way ANOVA with Tukey’s multiple comparison test or an unpaired t-test. *p < 0.05, **p<0.01, ***p<0.001. Differences between vehicle-treated WT mice and mice with CED and between vehicle- and ZA-treated mice with CED are presented.

### ZA restores the deteriorated osteocyte lacunocanalicular network in the cortical bone of mice with CED

While the increase in the bone volume of CED mice with ZA was traced back to reduced osteoclast number, the cellular mechanism contributing to increased bone strength in ZA treated CED mice was unclear. Previous studies have shown that changes in TGFβ signaling can profoundly impact osteocytes and their lacuno-canalicular network (LCN) in bone (Dole et al., 2017). Moreover, the intactness of LCN is often directly associated with higher bone strength (Dole et al., 2017, Dole et al., 2020, and Tang et al., 2013). Given this association between TGFβ signaling, osteocyte LCN integrity, and bone strength we examined the osteocytic LCN in this CED model. In the vehicle treated CED mice, that have exacerbated TGFβ signaling, we observed a 2.5-fold reduction in the LCN area compared to WT. This reduction in LCN area, is mostly likely a result of severe reduction in the length of canaliculi of the osteocyte network. With ZA treatment, the LCN area of WT mice remained unaffected, However, in comparison to the vehicle treated CED mice, ZA caused a 2-fold increase in LCN area of CED mice (Figure 5D, E). Together, these data suggest that excessive TGFβ signaling in CED mice leads to a severely deteriorated osteocyte LCN and that is returned to normal (i.e. similar to WT mice) with ZA treatment. Since reduced LCN correlates to reduced bone strength, it is likely that both the low bone strength and ZA induced improvement in bone strength of CED mice are in part attributed to changes that restore the osteocyte LCN.

Together, these data indicate that excessive TGFβ signaling in CED mice deregulates osteocyte mediated perilacunar/canalicular remodeling (PLR) which may contribute to reduced bone strength, which was prevented by inhibiting TGFβ release with ZA.

### 10 weeks of ZA treatment improves muscle function in CED mice

We measured longitudinal forelimb grip strength over 10 weeks to determine muscle function. Forelimb grip strength was significantly reduced in CED mice compared to WT mice, indicating muscle weakness in CED mice. Ten weeks of ZA-based treatment significantly increased grip strength in mice with CED but did not affect it in WT mice (Figure 6A). We next measured EDL whole muscle contractility (muscle specific force) and found that CED vehicle treated mice had reduced muscle contractility compared to WT mice treated with vehicle. In contrast with the 4 week treatment with ZA, after 10 weeks of ZA treatment, mice with CED exhibited significant improvements in muscle contractility (Figure 6B). Repeated tetanic stimulation of the EDL was performed to measure fatigability and we observed increased EDL fatigability in mice with CED compared to WT, which was improved with ZA treatment (Figure 6C). CED-Vehicle treated mice exhibited reduced TA muscle weight compared to WT and this was also improved with ZA treatment. There was no difference in EDL, gastrocnemius, or soleus weight between any of the groups (Figure S2). Notably however, ZA treatment did not affect muscle weight in WT mice (Figure S1D).

**Figure 6.**
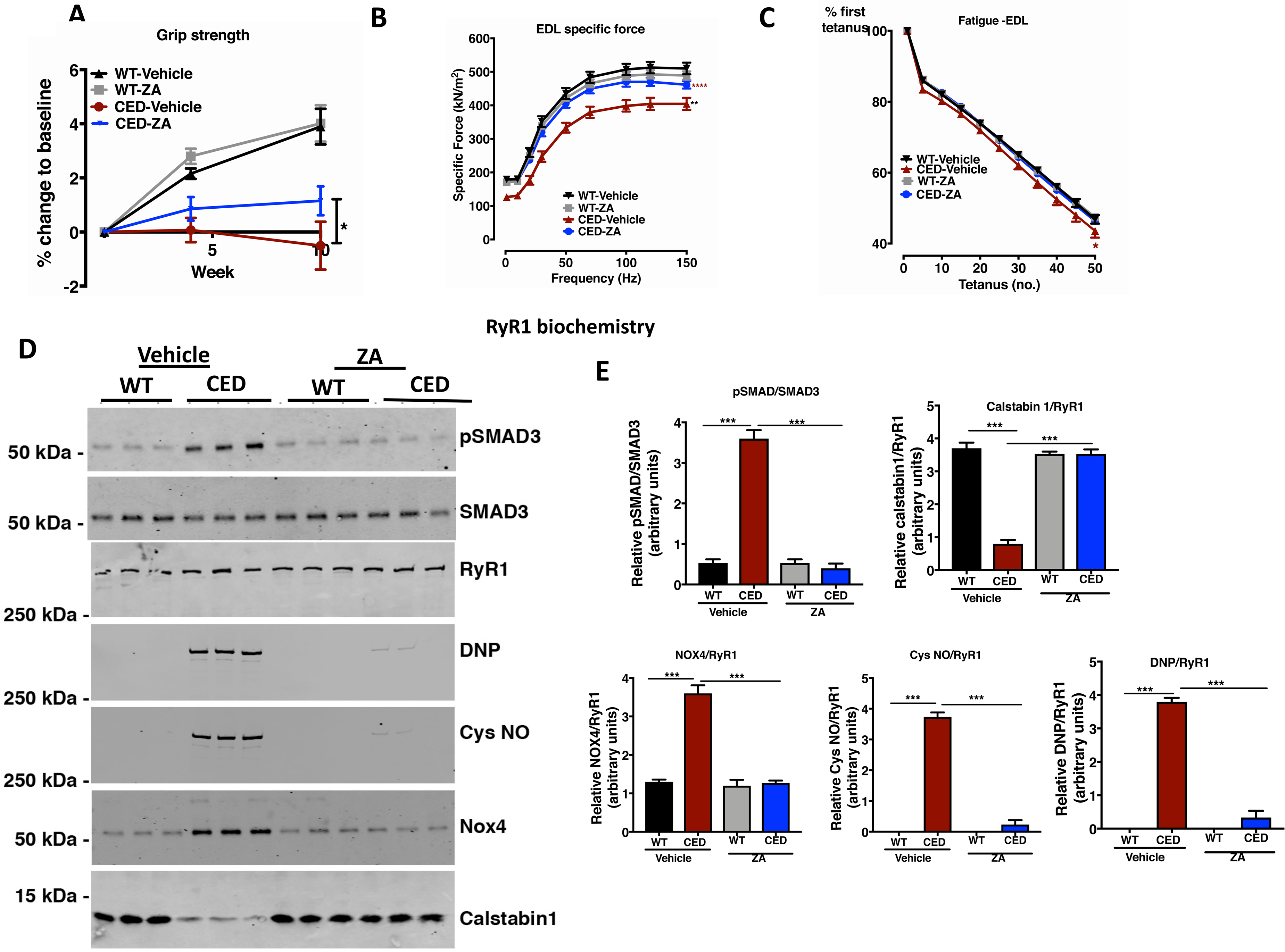
Skeletal muscle function (grip strength and *ex vivo* EDL muscle contractility), hindlimb muscle weights, and co-immunoprecipitation of RyR1, calstabin 1, and Nox4 oxidation and nitrosylation in EDL muscle. (A) Forelimb grip strength in mice was measured at baseline, week 5, and week 10 following ZA treatment. Changes in grip strength are expressed as percent change at baseline. (B) Muscle-specific force of the EDL muscle determined 10 weeks after ZA treatment. Muscle force was normalized by according to muscle weight and length. (C) EDL muscle fatigability measured after measurement of contractility. (D) Immunoblot of SMAD3 phosphorylation from EDL muscle showing co immunoprecipitation of Nox4 oxidation (DNP) and nitrosylation (Cys NO) and of RyR1 calstabin 1 binding. (E) Quantification of SMAD phosphorylation relative to SMAD3. Differences in the normalization of calstabin 1, Nox4, Cys NO, and DNP according to RyR1 are shown. Data are expressed as means (± SEM), and differences in grip strength, muscle function, and fatigue were determined using two-way ANOVA with Bonferroni’s multiple comparison test. *p < 0.05, **p<0.01, and ***p<0.001. Changes in muscle weight were determined using one-way ANOVA with Tukey’s multiple comparison test or an unpaired t-test. ***p<0.001. Differences between vehicle-treated WT mice and mice with CED and between vehicle- and ZA-treated mice with CED are presented.

### ZA improves muscle contractility in mice with CED by preventing Nox4-mediated oxidation of ryanodine receptor 1

Sarcoplasmic reticulum ryanodine receptor 1 (RyR1) is the major calcium release channel in skeletal muscle required for excitation-contraction coupling and regulation of muscle calcium homeostasis. Oxidation of RyR1 and loss of calstabin 1 binding to RyR1 in skeletal muscle lead to calcium release from the sarcoplasmic reticulum, which is associated with skeletal muscle weakness (Waning et al., 2015, Andersson et al., 2011, Bellinger et al., 2009). We have shown that increased TGFβ signaling in skeletal muscle increases Nox4 expression and Nox4-RyR1 interaction, which causes RyR1 oxidative stress and muscle dysfunction in mice. EDL muscle from CED mice had increased SMAD3 phosphorylation (pSMAD3), a downstream marker of TGFβ signaling. Consistent with TGFβ regulation of Nox4 expression, the muscles from these mice exhibited increased Nox4-RyR1 interactions and RyR1 oxidation and nitrosylation with loss of calstabin 1 binding to RyR1 (a biochemical signature of a RyR1 channel calcium leak). ZA treatment of CED mice reduced SMAD3 phosphorylation, reduced Nox4-RyR1 co-IP, reduced RyR1 oxidation and nitrosylation, and increased RyR1-calstabin 1 co-IP (Figure 6D, E). These data suggested that muscle weakness in mice with CED is due to Nox4-mediated oxidation of RyR1 resulting in calcium leak from the sarcoplasmic reticulum and that treatment with ZA improves muscle function by preventing the oxidation of RyR1.

### ZA improves muscle fiber structural properties in mice with CED

Similar to the increased phosphorylation of SMAD3 in the EDL muscle (Figure 6E), immunohistochemical staining of rectus femoris (RF) muscle samples from mice with CED showed an increase in SMAD2/3 phosphorylation. This effect was reduced by ZA-based treatment (Figures 7A and 7B), confirming that bone-derived TGFβ has systemic skeletal muscle effects. In addition, RF muscle fiber diameters and fiber cross-sectional areas were reduced in mice with CED. Treatment with ZA increased fiber diameters and cross-sectional areas as indicated by black lines in representative images of muscle sections (Figures 7A and 7C). Using *in vivo* micro-CT, the tibia was scanned to determine mid-calf muscle cross-sectional area. Mid calf muscle cross-sectional area was significantly reduced in CED mice relative to WT mice. ZA treatment showed increased mid-calf cross sectional area by micro-CT in CED mice, which was consistent with findings of immunohistological analysis (Figures 7D and 7E). In addition, we did not observe muscle fibrosis in mice with CED with or without this treatment ZA treatment (Figure S3).

**Figure 7.**
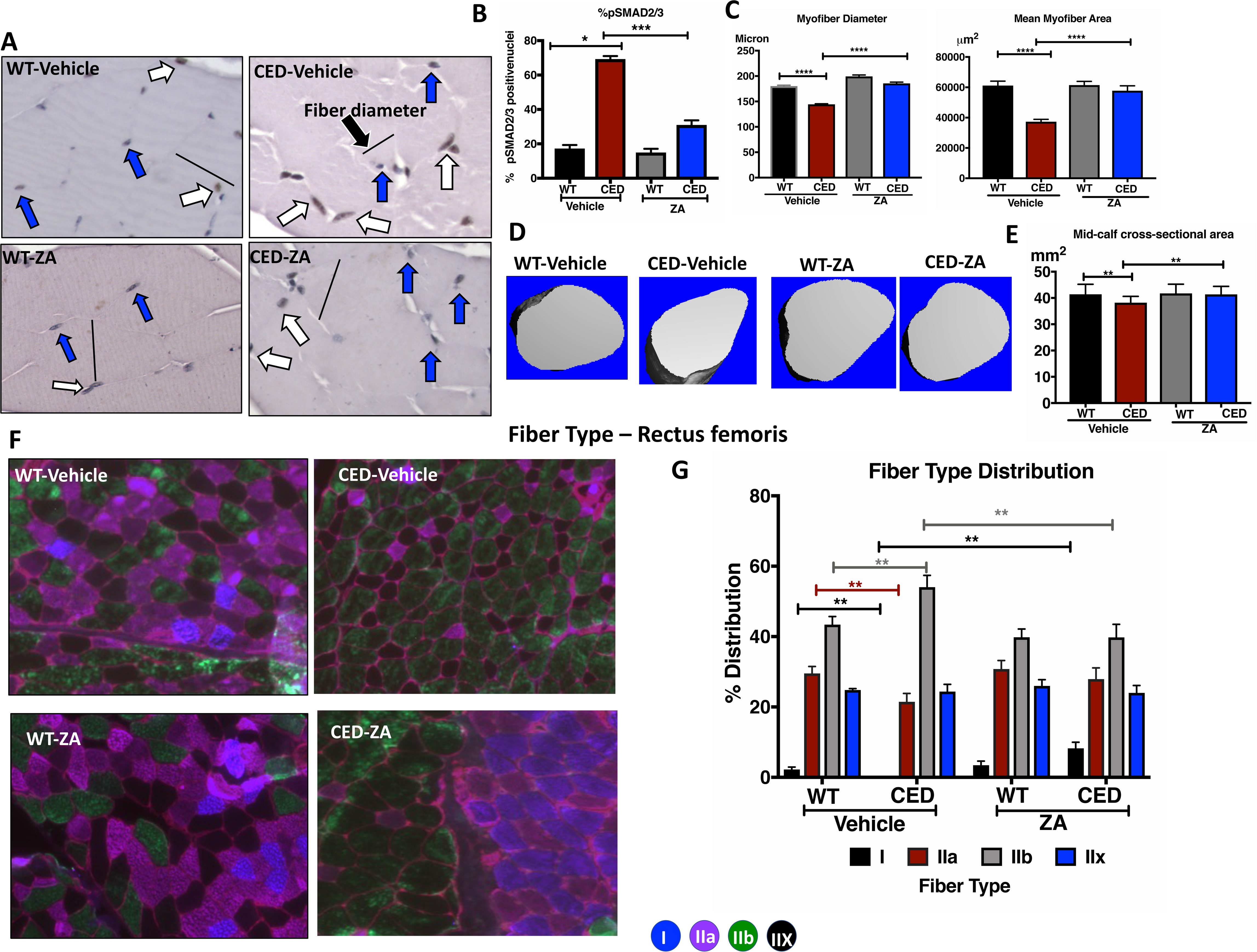
Skeletal muscle histology, immunohistochemistry, micro-CT, and immunofluorescent myosin heavy chain fiber typing using a cocktail antibody method. (A) Representative images of muscle sections stained with an anti-pSMAD2/3 antibody. The blue arrows indicate pSMAD2/3-negative cells, the white arrows indicate brown-colored pSMAD2/3-positive cells, and the black lines indicate fiber diameters. (B) Quantitation of pSMAD2/3 expression in paraffin-embedded RF muscle sections using immunohistochemistry. The ratio of pSMAD2/3-positive nuclei to total nuclei is presented as a percentage. (C) Myofiber diameters and muscle cross-sectional areas in RF muscle sections measured using BIOQUANT image analysis software. (D) Representative micro-CT reconstructions of mid-calf muscle cross-sectional areas (E) Mid-calf muscle cross-sectional areas assessed using in vivo micro-CT scans of proximal tibiae. (F) Frozen RF muscle sections processed using isopentane and used for immunofluorescence with specific antibodies against type I, type IIa, and type IIb myosin heavy chains. Slow-twitch fibers are shown in blue, intermediate fibers (between type I and IIa) are shown in purple, fast twitch fibers are shown in green, and intermediate fibers (between type IIa and IIb) are shown in black. All fibers were separated using laminin immunostaining. (G) Differences in quantification of fiber types presented as a percent distribution. In all groups, all type II fibers were significantly higher than type I fibers. Data are expressed as means (± SEM), and differences between groups were determined using one-way ANOVA with Tukey’s multiple comparison test or an unpaired t-test. *p < 0.05, **p<0.01, ***p<0.001, and ****p<0.0001. Differences between vehicle-treated WT mice and mice with CED and between vehicle- and ZA-treated mice with CED are presented.

We also determined the distribution of muscle fiber types in mice with CED and the possible effect of ZA on fiber type shifting. RF muscles from CED mice exhibited reduced oxidative type I and IIa fibers (slow twitch fibers) and an increase in glycolytic type IIb fibers (fast twitch fibers) compared to muscles from WT mice. ZA treatment significantly increased the percentage of type I fibers and reduced that of type IIb fibers in mice with CED (Figures 7F and 7G). Similarly, the TA muscle of CED mice also had fewer slow twitch (type IIa) and greater fast twitch (type IIx) and with ZA treatment the fiber type shifted back from type IIx to IIa (Figure S4A). The reduction in the percentage of slow twitch fibers observed in CED mice is similar to the reduced slow twitch fibers observed in many muscle diseases such as myotonic dystrophy type I, and other condition such as obesity, diabetes, prolonged bed rest and in mouse models of disuse. The increased fast twitch type IIb fibers in the RF muscle and type IIx fibers in the TA muscle might have been associated with the increased rate of fatigue in CED mice (Figure 6C). ZA treatment in WT mice did not have any effect on fiber type distribution, which suggests ZA is effective in restoring normal muscle fiber type distribution specifically in diseased states.

### ZA increases mobility in mice with CED

Open field test has been used to assess locomotion and general activity in mice (Seibenhener and Wooten, 2015). In our study, vehicle-treated mice with CED exhibited a trend of reduced mobility as indicated by fewer entries into the center and periphery of an activity monitor when compared with vehicle-treated WT mice (Figure S4). Consistent with these data, CED vehicle treated mice had reduced movement in the plus elevated maze, spent less time in open arm of the maze and tended to remain in one corner of the maze. ZA-based treatment of CED increased the mice’s mobility in both open field and elevated plus maze (Figure S4B). Because behavioral studies require relatively large numbers of mice to achieve appropriate power, we did not observe significant changes in the mobility detection results of these tests. However, the trend strongly favors reduced mobility in response to increased bone turnover and muscle weakness in mice with CED, which treatment with ZA overcame.

### ZA has potential direct effects on skeletal muscle mass and function

Our *in vivo* results showed that ZA treatment was associated with improved muscle function by blocking bone resorption and preventing TGFβ release from bone in a high bone turnover state. To determine whether ZA has direct effects on myoblasts in the presence or absence of TGFβ, we used *in vitro* cell culture assays to determine the proliferation and differentiation potential of C2C12 myoblasts with and without ZA treatment (Figure S5). Consistent with previous findings, TGFβ induced the proliferation of these cells (Krueger and Hoffmann, 2010, Wicik et al., 2010,

Delaney et al., 2017). In the presence of TGFβ, ZA doses of 0.625 and 1.25 mM reduced C2C12 cell proliferation and induced their differentiation (Figures S5A and S5B). Higher doses of ZA (≥50 mM) were lethal to C2C12 cells in the presence of TGFβ (Figure S2). Therefore, in subsequent experiments with C2C12 cells, we used ZA doses of 0.625 and 1.25 mM.

We also determined the differentiation potential of C2C12 myotubes in the presence and absence of TGFβ and ZA. We used the fusion index (the ratio of the number of nuclei inside a myotube to that outside of it), number of myotubes, and average number of nuclei per myotube to determine the potential of C2C12 differentiation. Treatment with TGFβ significantly lowered the fusion index, reduced myotube numbers, and resulted in fewer average nuclei per myotube compared to vehicle treatment (Figures S5B and S5C). Adding ZA to TGFβ-treated cultures of C2C12 myotubes significantly increased the fusion index, number of myotubes, and average number of nuclei per myotube, restoring their levels to those in untreated C2C12 cells (Figure S2). TGFβ treatment also reduced myotube diameters (Figure S5B), which was consistent with the reduced muscle fiber diameters observed in mice with CED *in vivo* (Figure 7C). In the presence of TGFβ, ZA significantly improved the differentiation of C2C12 myotubes (Figures S5B and S5C). Taken together, these data showed that in the presence of TGFβ, ZA exerts a direct effect on myoblasts, inhibiting TGFβ-induced proliferation and promoting differentiation. Also, TGFβ-based treatment of C2C12 cells increased SMAD3 phosphorylation, increased RyR1 oxidation and nitrosylation, and reduced binding of calstabin 1 to RyR1 (Figure S5D). Adding ZA to C2C12 cultures containing TGFβ prevented TGFβ-induced Nox4-mediated oxidation and destabilization of RyR1-calstabin 1. Collectively, these *in vitro* data showed that excess TGFβ inhibits myotube differentiation, whereas ZA-based treatment promotes the differentiation of myoblasts. Notably, ZA did not have negative effects on muscle either *in vitro* or *in vivo* (Figures S1 and S5B).

## DISCUSSION

Pathological loss of bone results in loss of muscle mass and function, leading to debilitating consequences like falls, fractures, and disability (Cianferotti and Brandi, 2014). The present study advanced our understanding of bone to muscle signaling. Several nonmalignant bone disorders are associated with muscle weakness (Beaudart et al., 2018, Gentry et al., 2010), and in this study, we used a mouse model of high bone turnover (CED) with excess TGFβ and identified that the U.S. Food and Drug Administration-approved antiresorptive drug ZA (Dhillon, 2016, Rakel et al., 2011, Brown and Guise, 2009, Coleman et al., 2011) improves both bone and muscle function.

CED is a metabolic bone disease caused by constitutively active TGFβ, resulting in extremely high bone turnover and incidence of fractures and deformities (Tang et al., 2009, Janssens et al., 2006). Mice with CED exhibit increased bone density at the diaphysis similar to that seen in CED patients (Bondestam et al., 2007). Long-term ZA administration in children and young adults with CED improves their bone lesions, normalizes their levels of bone turnover markers, and increases their BMD (Baroncelli et al., 2017). Similar to CED patients, in the mouse model of CED, we found that both short-term (4 weeks) and long-term (10 weeks) ZA use increased BMD and improved the microarchitecture in the long bones and vertebrae. In a 10 week treatment experiment, longitudinal dual-energy x-ray absorptiometry (DXA) did not show any differences in BMD between WT mice and mice with CED, probably due to the two dimensional and nonvolumetric nature of DXA. However, small volumetric changes in bone microarchitecture can be quantified using high-resolution micro-CT (Genant and Jiang, 2006). In the present study, micro-CT showed a significant reduction in trabecular bone volume and increased cortical bone porosity in CED mice. Also, CED patients have increased cortical thickness as determined via radiography. However, our data using details micro-CT analysis of bones demonstrated that cortical thickness is significantly reduced in mice with CED. Our results suggest that, besides radiography and DXA, introduction of micro-CT in characterizing CED in patients can improve diagnosis and monitoring of treatment response.

Improvement in bone mass itself is not sufficient to improve bone strength (Schuit et al., 2004, Sornay-Rendu et al., 2007). Bone volume and material quality (Schuit et al., 2004, Sornay Rendu et al., 2007) as well as muscle mass and function (Amthor et al., 2007) are necessary for tissues to function well both individually and in coordination (Kim and Kang, 2017, Wagner et al., 2018, Schuit et al., 2004). Despite increased BMD, the bones of CED patients have low bone material strength indices (Herrera et al., 2017). Osteocyte-mediated remodeling of the perilacunar/canalicular matrix is essential for bone quality (Dole et al., 2017). TGFβ coordinates the functions of osteoblasts, osteoclasts, and osteocytes, thus maintaining bone mass and quality (Mundy, 1991, Edwards et al., 2010, Mohammad et al., 2009). In CED, excess TGFβ deregulated osteocyte-mediated perilacunar/canalicular remodeling, which may be a contributing factor for reduced mechanical properties in mice with CED. Absence of TGFβ signaling in bone causes defects in perilacunar/canalicular remodeling (Dole et al., 2017), and our data add that excess TGFβ signaling in bone also causes deregulation of osteocyte-mediated bone remodeling and associated fragility, suggesting that optimal TGFβ levels are required for normal bone homeostasis. We observed that with ZA treatment, mice with CED have reduced cortical porosity and increased resistance to fractures. Treatment with ZA normalized serum TGFβ levels and rescued the deregulation of PLR in CED mice, which in turn associated with increased mechanical strength of the bone from mice with CED. Authors did not report the effect of ZA in improving cortical bone quality in an earlier study in CED patients (Baroncelli et al., 2017). With radiographic assessment, separating ZA treatment-mediated bone formation from sclerosis at the diaphysis of tibia due to CED is difficult (Janssens et al., 2006, Tang et al., 2009). We also observed that in addition to improving both bone volume and muscle mass, ZA can improve bone mechanical properties. Thus, improved bone microarchitecture and mechanical integrity with ZA are indicators of fracture resistance in an extremely high bone turnover state.

TGFβ is a known stimulator of osteoblastic bone formation (Mundy, 1991, Bonewald and Mundy, 1990, Wu et al., 2016). In mice with CED, we found that excess TGFβ signaling increased the number of osteoclasts and osteoblasts and promoted the bone formation rate, a typical characteristic of CED mouse models. In a previous study (Tang et al., 2009), investigators determined the numbers of osteoclasts and osteoblasts at the endosteal surface and found that they were increased in mice with CED. In the present study, we observed similar changes in bone cells but at the trabecular bone surface, the most reactive bone site; this finding adds further details to the characterization of CED. TGFβ has a dual effect on osteoclastogenesis, which is largely dependent on the stage of osteoclast differentiation (Quinn et al., 2001, Tang and Alliston, 2013). We also observed increased osteoclast activity in mice with CED. Despite the increased bone formation rate, CED mice have an overall reduction in trabecular bone volume. Blockade of TGFβ signaling using an anti-TGFβ antibody or TGFβ receptor inhibition has increased bone mass and quality in normal mice (Mohammad et al., 2009, Edwards et al., 2010). Suppression of both osteoclast and osteoblast numbers along with reduced bone formation rates in mice with CED after ZA treatment suggests that either the entire bone remodeling process is being suppressed or that the bone in under healing conditions with repair of the osteocyte lacunocanalicular network and increasing bone material strength and volume (Huja et al., 2009). Collectively, these data suggest that in a TGFβ-induced extreme bone turnover state, ZA-induced inhibition of TGFβ release from bone improves both bone mass and quality by normalizing the coordination of osteoblasts, osteoclasts, and osteocytes.

We next identified whether loss of bone mass and strength caused by excess TGFβ can impact muscle function in mice with CED. About 39% of CED patients experience muscle weakness, whereas 21% have reduced muscle mass (Janssens et al., 2006) due to unknown mechanisms. Also unclear is whether muscle weakness in CED and those with abnormal bone turnover is a result of loss of muscle mass, muscle strength, or both. Detecting the initiation of muscle weakness in patients with bone disorders is important for early therapeutic intervention. In our study of mice with CED, although bone destruction was evident at birth due to mutation of the TGFβ gene, muscle weakness only became evident after 8 weeks of age. The delayed onset of muscle weakness may have been due to the necessity of sustained exposure to high TGFβ in the bone-muscle milieu. In contrast with bone metastases, bone loss is a primary event in CED cases, and if untreated, it could cause continuous release of bone-derived TGFβ and muscle weakness. Thus, treatment of bone loss can concurrently improve muscle function by blocking the release of matrix-derived factors. Previously, in mice with bone metastases, we showed that muscle dysfunction can appear before loss of muscle mass (Waning et al., 2015). In the present study, we observed that improvement of muscle function takes longer than that of muscle mass, as prolonged ZA treatment was required to improve muscle function. We observed improvement in bone mass and muscle fiber diameter in mice subjected to 4 weeks of ZA administration, but improvement in muscle function only in those subjected to 10 weeks of ZA. Improvement in muscle function was associated with stabilization of RyR1 calcium channels and reduction of oxidative stress, which increased muscle contractility. Short periods of ZA treatment may not be sufficient to reduce TGFβ to that allow improvement in muscle strength. In bone diseases with continuous bone loss, the bone and muscle environments are under continuous exposure to TGFβ. Under these conditions, long ZA exposure may be required to normalize TGFβ-mediated downstream effects such as muscle weakness.

In patients with bone diseases and associated muscle weakness, ZA exerts broad beneficial effects on muscle by inhibiting the constant release of active TGFβ from bone, thus improving the muscle phenotype and preventing muscle dysfunction. Researchers have shown that myocyte hypertrophy may not translate into increased muscle strength (Amthor et al., 2007) because tendons, connective tissues, and energy metabolism must also adapt in response to exposure to drugs or other hypertrophic stimuli (Laurent et al., 2016). In contrast with the 10 week treatment in our study, shorter treatment with ZA may not have impacted these factors and thus may have resulted in a lack of improvement in muscle strength in mice with CED.

ZA has been used as a prophylactic therapy in patients with high risk of bone loss such as post-menopausal osteoporosis, cancer or cancer treatment associated bone loss (Dhillon, 2016, Santa-Maria et al., 2018, Sun et al., 2016, Ottanelli, 2015, Ganguly et al., 2012, Boissier et al., 2000, Hornby et al., 2003). In the present study, ZA-based treatment in mice with CED prior to apparent muscle weakness prevented the development of muscle weakness, suggesting that a preventive ZA use strategy has beneficial effects on reducing muscle weakness in patients who are at high risk for musculoskeletal disease. Our data demonstrating that the use of ZA as a preventive treatment of muscle weakness are supported by reports that adjuvant ZA-based treatment with chemotherapy prevented muscle weakness in mice (Hain et al., 2019, Essex et al., 2019). Furthermore, activation of TGFβ signaling contributes to pathological fibrosis in skeletal muscle, impairing the force-generating capacity of the muscle (Ismaeel et al., 2019). However, we did not observe any muscle fibrosis in mice with CED, indicating that muscle weakness in these mice is primarily due to abnormal bone pathology, not to changes in muscle tissue. Absence of muscle toxicity and fibrosis with ZA treatment in WT mice suggests that use of ZA as a preventative medicine for muscle weakness is a safe strategy.

Mice with CED have extreme bone abnormalities, especially at the tibial diaphysis, due to the formation of callus, fibrosis, and fractures. In the present study, the TA muscle, which lies in close proximity to this region, had necrotic areas in the mice with CED which may have reduced the TA muscle weights. Due to the close physical attachment of bone and muscle, the callus and fractures in bone may have interrupted muscle vasculature, leading to TA muscle damage, which was reflected in macroscopic necrosis and muscle damage (Davis et al., 2015). ZA-based treatment resulted in increased TA muscle weight, possibly due to improvement in bone loss and fractures, leading to reduced muscle damage. In our future studies, we will determine underlying mechanisms of fracture-associated disruption of muscle vasculature in cases of bone diseases. We also found that reduced mid-calf muscle cross-sectional area, RF muscle myofiber area and diameter, forelimb grip strength, and EDL muscle contractility in a hindlimb in mice with CED improved in response to ZA use, indicating that the effects of ZA are not limited to a single muscle type.

Muscle fiber types switching/selection has a profound impact on muscle diseases, which makes this process an important therapeutic targets in the prevention and treatment of metabolic diseases (Talbot and Maves, 2016). Muscles have varying proportions of slow-twitch (type I) and fast-twitch (type II) fibers depending on their functional requirements. Slow-twitch fibers are oxidative and resistant to fatigue, whereas fast-twitch fibers are glycolytic and easily fatigable (Grimby et al., 1976). During muscle loss due to disuse, heart failure, spinal cord injury, or chronic obstructive pulmonary disease, fiber types shift from type I to type II (Talbot and Maves, 2016, Trappe et al., 2007, Baldwin et al., 2013). In our study in CED mice, we observed the shift from type I to type II fibers in the RF and TA muscles as well as increased EDL muscle fatigability, which indicates muscle atrophy. Taken together our data suggest that in CED mice severe bone and muscle deformities and associated pain may have restricted their mobility, as seen in open field and elevated plus maze tests. Such muscle adaptations may be essential for movements that are fast or quick and thus essential to survival rather than movements requiring sustained endurance. In human studies and tests using rodent models, mechanical unloading and disuse reduced muscle power and induced slow-twitch to fast-twitch fiber shift, which was reversed by exercise and pharmacological interventions, possibly through increased neuromuscular activation (Gallagher et al., 2005, Trappe et al., 2007, Takemura et al., 2017, Sharlo et al., 2019, Lawler et al., 2014, Baldwin et al., 2013, Desaphy et al., 2010). Consistent with data from studies of humans and rodent models, increased movement in mice with CED in response to ZA administration correlates with a reversion of fiber type shift from fast to slow together with reduced muscle fatigue and increased mobility in open field and elevated plus maze tests. Thus, extreme bone turnover mediated by TGFβ shifts the muscle fiber type, and preventing TGFβ release and bone loss by ZA treatment reverses this shift to the normal state. These findings may lead to evaluation of the clinical utility of ZA in treating muscle weakness by shifting the fibers in patients with bone diseases, as restricted mobility is the major contributing factor that limits their day-to-day activity.

Consistent with previous reports, we observed that TGFβ inhibited myogenesis (Ugarte and Brandan, 2006, Zhu et al., 2004, Krueger and Hoffmann, 2010, Wicik et al., 2010, Hesse et al., 2019) and increased C2C12 cell proliferation (Krueger and Hoffmann, 2010, Wicik et al., 2010, Delaney et al., 2017). Adding ZA to TGFβ-treated cultures of C2C12 myotubes reduced the proliferation and induced myotube differentiation of C2C12 cells in the presence of TGFβ. These effects of ZA can be beneficial in patients with muscle injury and damage, in whom excess TGFβ delays muscle recovery and repair (Kim and Lee, 2017, Delaney et al., 2017, Hesse et al., 2019), as the addition of ZA to a treatment regimen can divert muscle toward differentiation without causing hypertrophy. This finding is consistent with a report that C2C12 myotube size increased when treated with serum from patients given bisphosphonates (Klein, 2020). Our *in vitro* data on C2C12 cells demonstrate the potential direct effects of ZA on skeletal muscle in the presence of TGFβ. If tested *in vivo*, this observation could have a broader impact on treatment of not only bone-associated muscle dysfunction but also non-bone-associated muscle diseases. Ascertaining whether ZA has a direct effect on muscles *in vivo* is challenging, as bone and muscle are tightly connected tissues, and the contribution of bone-mediated effects of ZA on muscle health cannot be excluded. Cell intrinsic mechanisms of ZA action on osteoclasts differentiation involve inhibition of mitogen-activated protein kinases (MAPK) and NF-κB pathways(Wang et al., 2022). Since both MAPK and its downstream signaling pathways are major contributors of regulation of many cell processes including cell differentiation, our future studies will determine potential effects of ZA in regulation of these pathways in differential of myocytes.

In patients with bone metastases, ZA is administered with a long dosing interval, which prevents osteolysis but does not reduce the skeletal tumor burden. However, investigators found that administration of ZA at a low dose on a daily or weekly basis reduced bone metastasis (Daubine et al., 2007). Several *in vitro* and *in vivo* studies using rodent models also showed that ZA has antitumor effects against bone and soft tissue metastases (Boissier et al., 2000).

Continuous or frequent intermittent low-dose ZA administration results in prolonged exposure of the bone microenvironment to ZA, thus enabling direct effects on tumor cells that reside in bone (Boissier et al., 2000, Daubine et al., 2007). Early cancer studies and data from the present study demonstrated that prolonged treatment with ZA is needed to reduce excess TGFβ levels. *In vivo* studies of prolonged exposure to low-dose ZA versus one high dose of ZA in mouse models of nontumor bone destruction with high TGFβ release would provide better insight into the direct effects of ZA on muscle. Studies in a setting of non-bone-mediated muscle weakness, such as cancer cachexia or age-associated muscular dystrophy, will also provide insight into the direct *in vivo* effects of ZA on improvement of muscle weakness.

### Limitations of the study

CED is an extreme case of TGFβ-induced bone turnover that might not be observed with many nontumor bone diseases (Wang et al., 2018, Tang et al., 2009). Abnormally high active TGFβ levels cause uncoupling of bone resorption and formation, a characteristic of rare genetic skeletal diseases such as CED, osteogenesis imperfecta, and other common musculoskeletal disorders, including osteoarthritis (Grafe et al., 2014, Zhen et al., 2013, Crane and Cao, 2014, Wang et al., 2018). Under these conditions, blocking bone resorption is a relatively complex process. Thus, in our CED mouse model, ZA treatment did not effectively inhibit bone resorption. Whereas CED is an extreme case, a useful direction would be to evaluate the muscle weakness in cases of other bone diseases in which TGFβ may be a central factor and determine whether the inhibition of TGFβ can improve muscle function. To that end, similar studies using mouse models of bone specific overexpression of TGFβ would provide better insight into bone to muscle signaling. Further evaluation of an anti-TGFβ antibody as a treatment strategy would be useful in understanding the bone to muscle signaling mediated by TGFβ. Recent studies showed that osteocytes can stimulate myogenesis and enhance muscle function (Huang et al., 2017), so whether TGFβ can directly regulate the osteocyte network to improve muscle function in a high bone turnover state must be determined.

### Conclusions

We determined the broad function of ZA in condition of skeletal muscle weakness using *in vitro* and in *vivo* models and a pharmacological treatment approach similar to those used in clinical trials for the treatment of common bone diseases. Using this approach, we identified the extended role of ZA in bone-induced muscle signaling through inhibiting the release of TGFβ from the bone matrix under non-tumor-associated bone conditions.

Our mouse model data on high bone remodeling suggest that systemic release of bone derived TGFβ caused dysregulation of the osteocyte network, promoted negative adaptation of muscle fibers, reduced muscle function and mass, and may have compromised mobility. These findings will have wide implications for muscle dysfunction in patients with metabolic bone diseases. The preventive use of ZA over a long period normalized TGFβ concentrations, rescued bone and muscle mass and mechanical properties, and improved overall musculoskeletal health. ZA-based treatment may heal bone destruction and simultaneously improve muscle mass and function in patients with tumor- and non-tumor-associated bone diseases. Such a strategy may also be beneficial for elderly patients who suffer from muscle weakness and bone loss, as bone destruction starts before muscle weakness, and preventing bone loss will improve muscle function, thus reducing the frailness and falls commonly observed in this population.

## ACKNOWLEDGMENTS

The authors thank Dr. Xu Cao for providing mice with CED. The authors also acknowledge Emily Pemberton for her assistance with the biomechanical testing of bones. We thank Don Norwood, Scientific Editor, Research Medical Library, for assistance in editing and formatting this article. Dr. Guise is a CPRIT Scholar in Cancer Research.

## AUTHOR CONTRIBUTIONS

T.T., D.L.W., T.A., K.S.M., and T.A.G. conceptualized the project. T.T., M.M., S.J., S.R, G.M.P., N.S.D., S.S., B.A.H., J.R., L.W., Y.S.,R.O., A.R., K.S.M, T.A.G performed the experiments and analyzed the data. T.T. and T.A.G. wrote the original draft of the manuscript. T.T., S.S. S.R., G.M.P., N.S.D., B.A.H., J.R., A.R.M, D.L.W., T.A., K.S.M., T.A.G. reviewed and edited the manuscript.

## STAR METHODS

### KEY RESOURCES TABLE

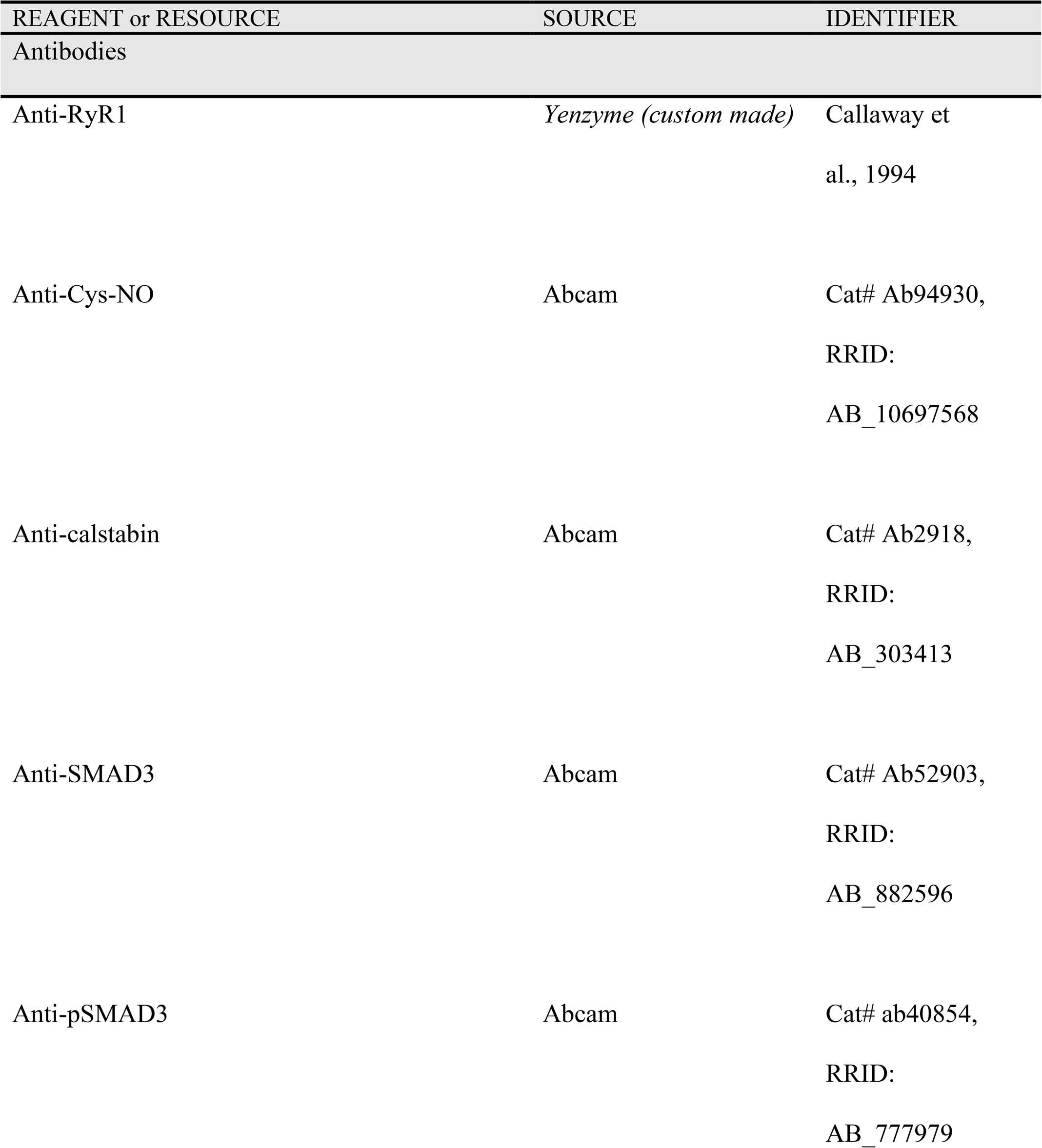
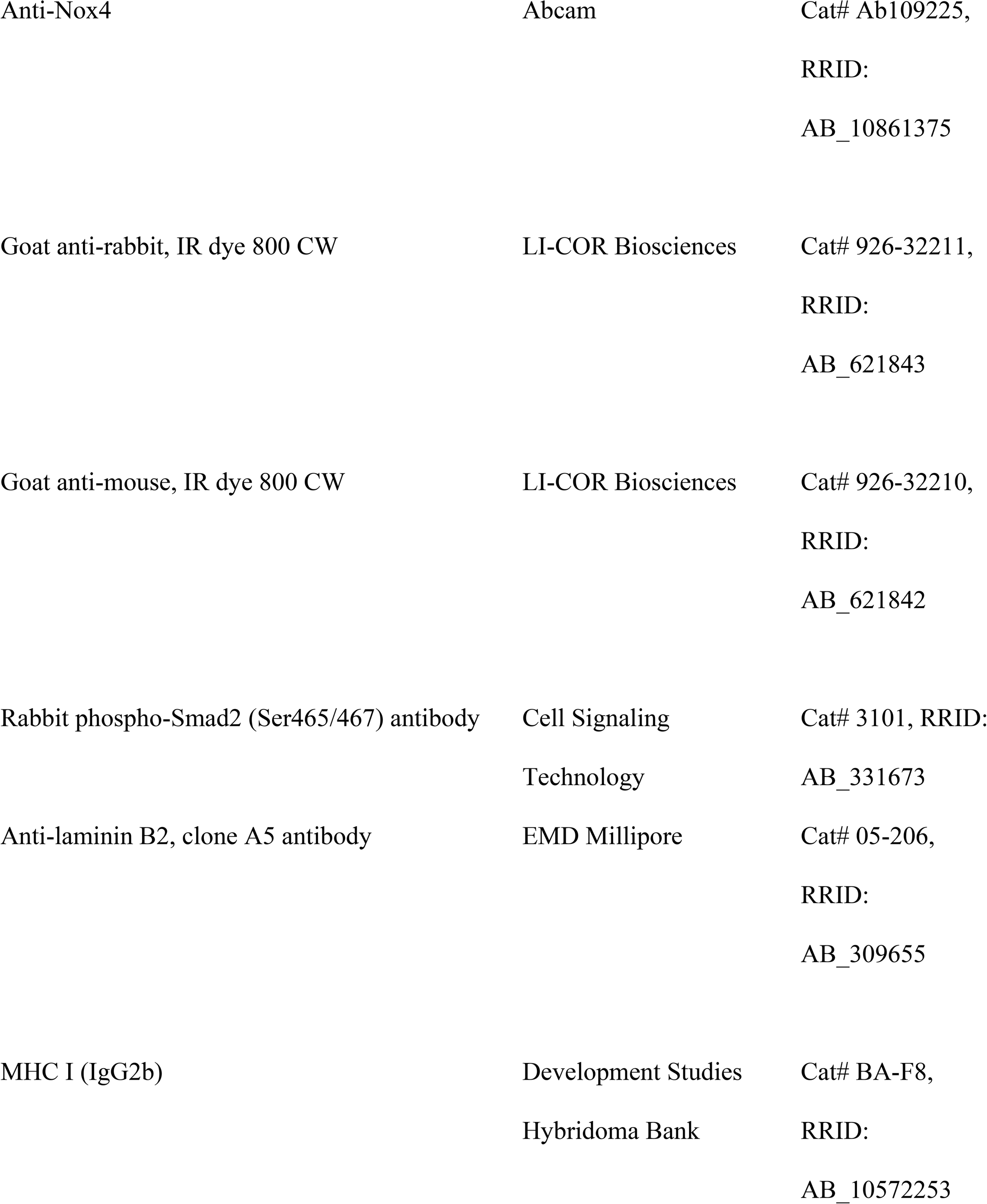
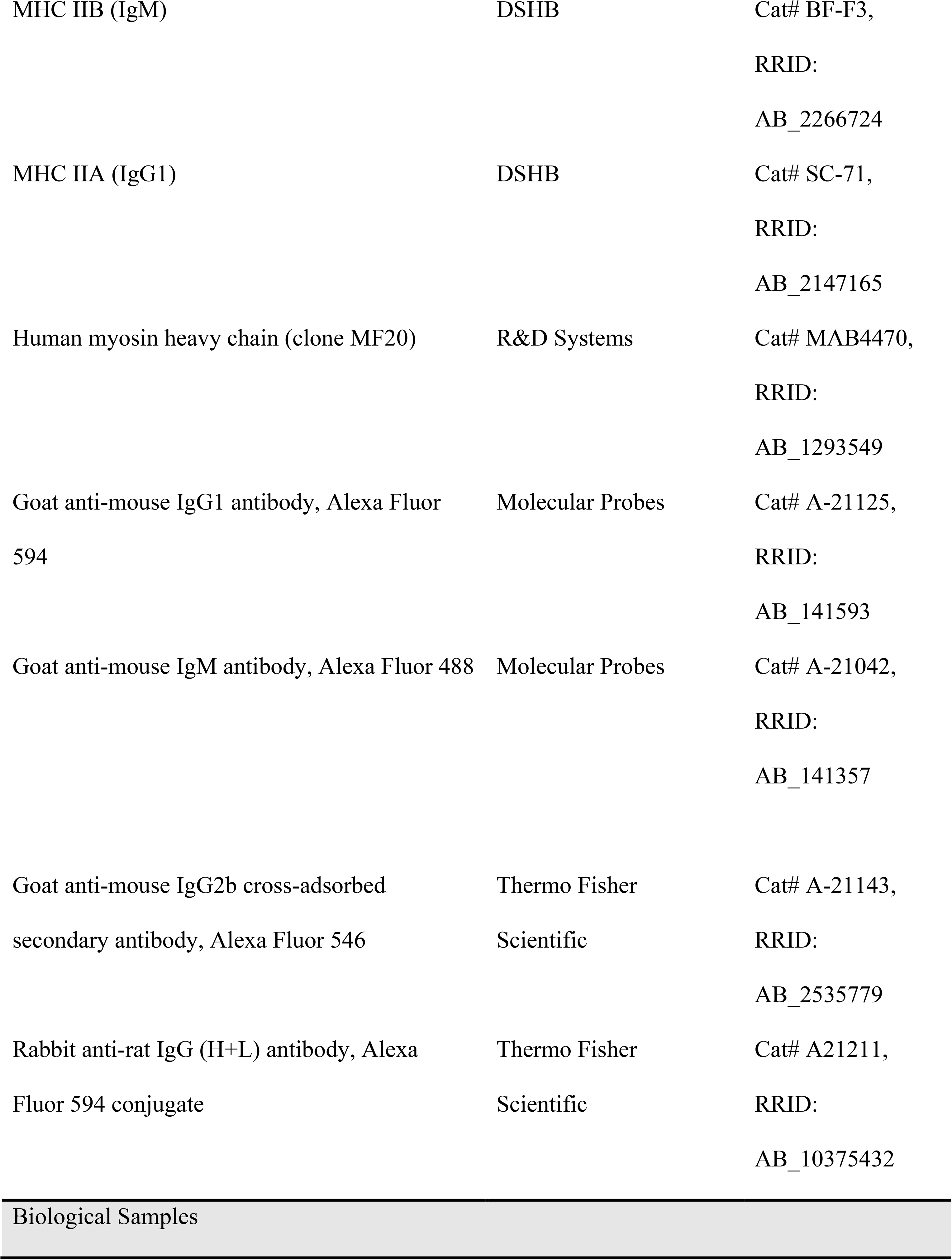
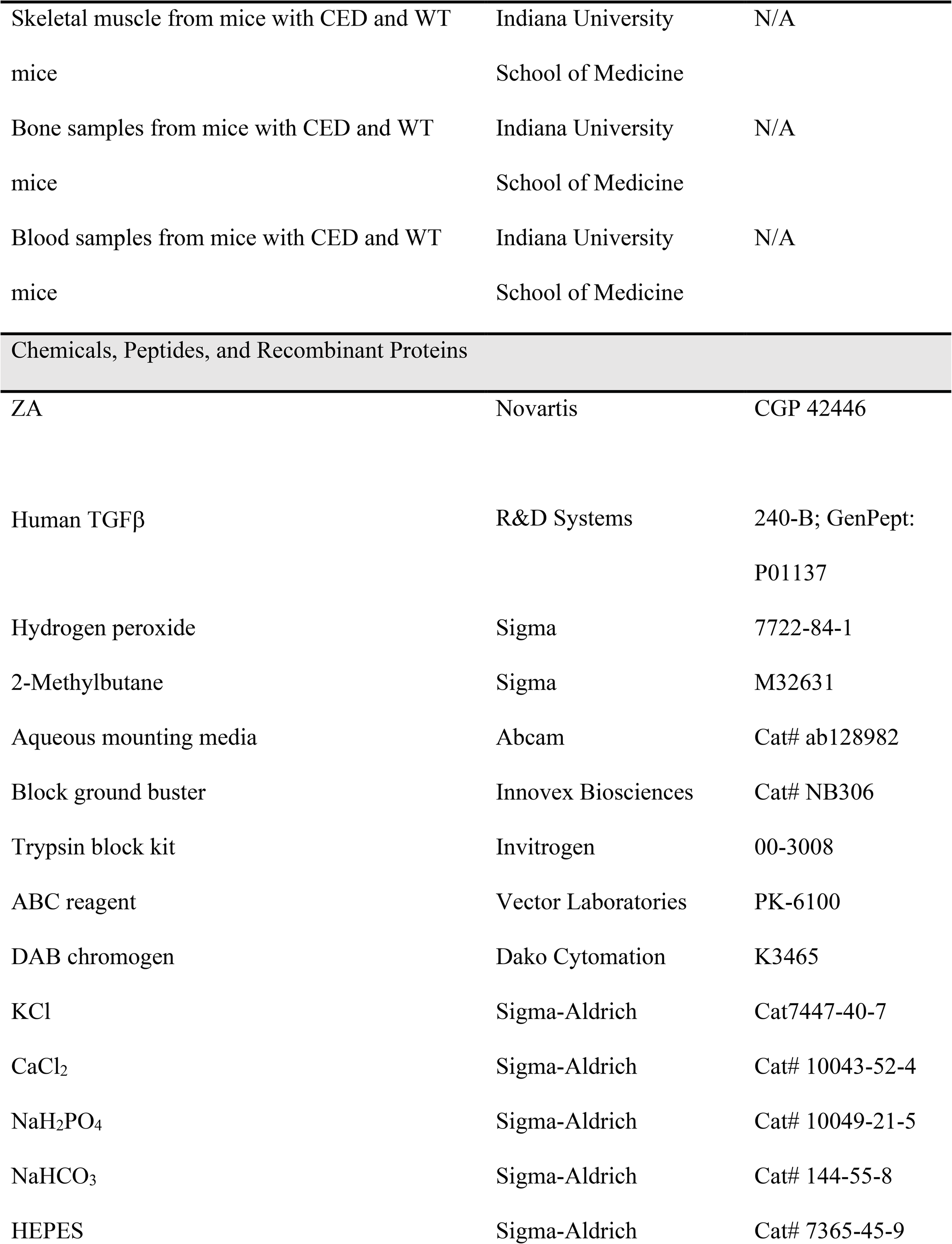
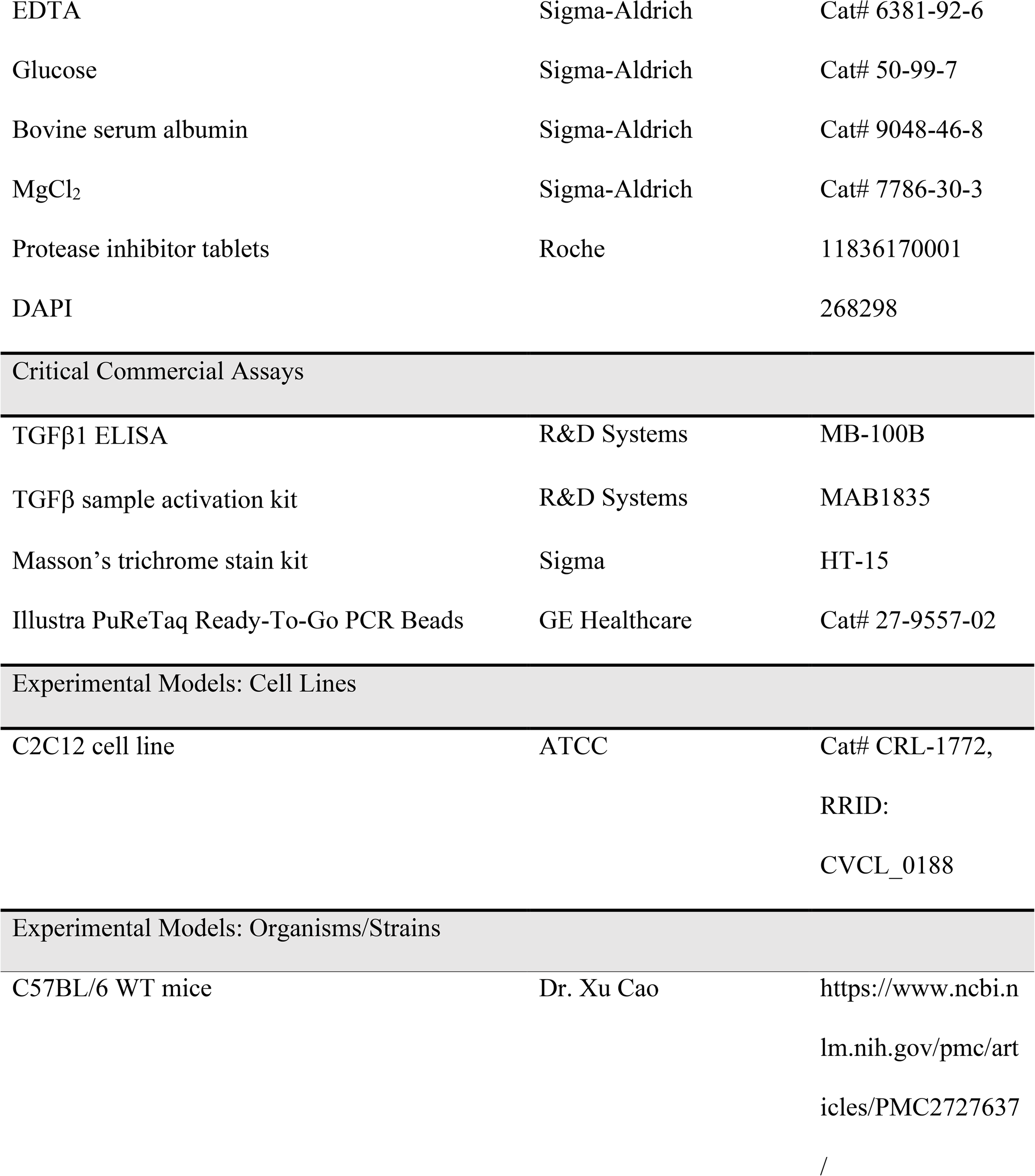
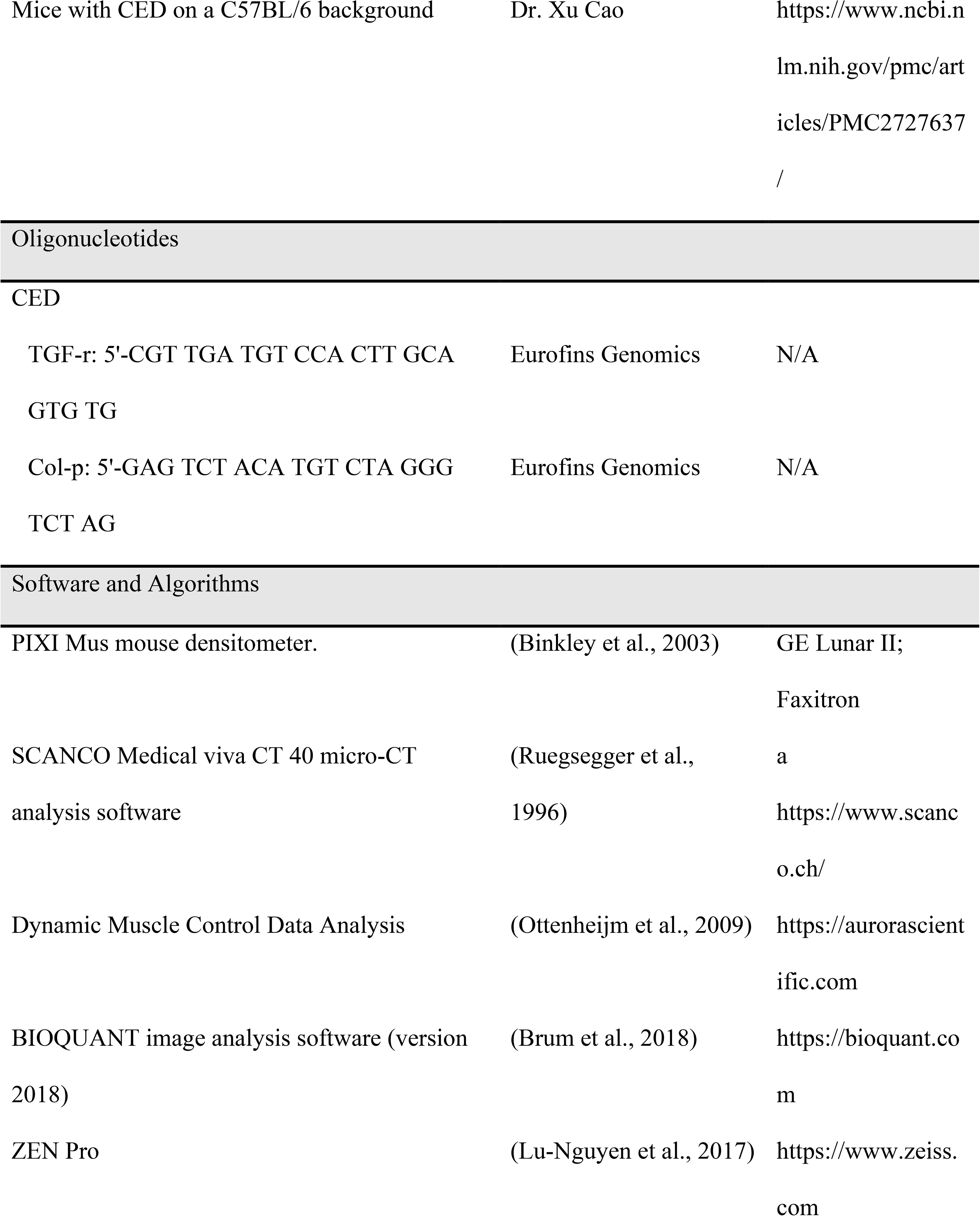
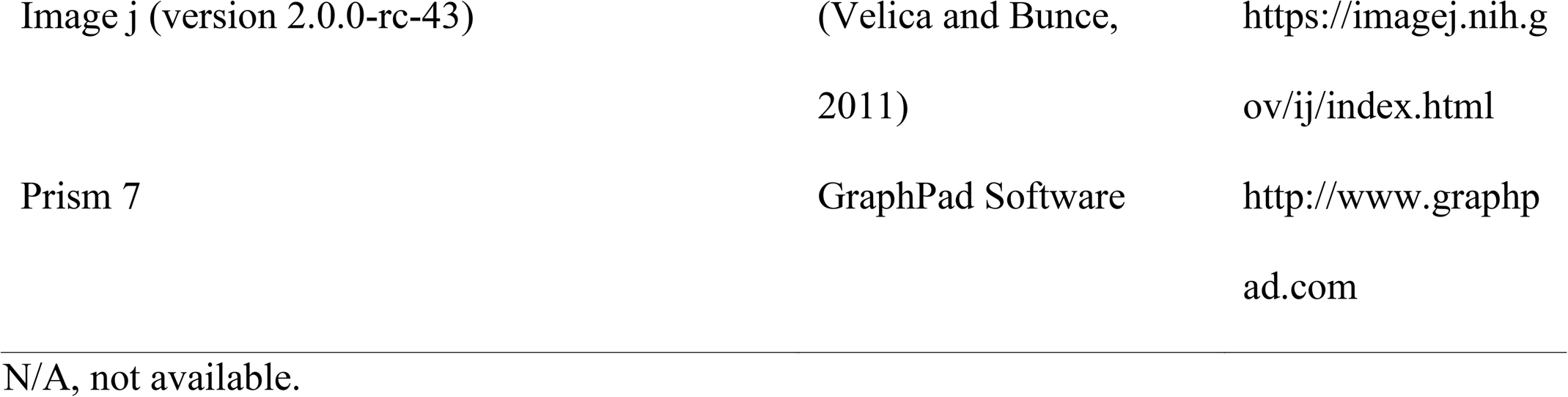

## CONTACT FOR REAGENT AND RESOURCE SHARING

Further information and requests for resources and reagents should be directed to and will be fulfilled by the lead contact, Trupti Trivedi.

## EXPERIMENTAL MODEL AND SUBJECT DETAILS

C57BL/6 mice with CED carrying a CED-derived *TGFβ1* mutation and their littermates WT for full-length *TGF-β1* (Tang et al., 2009) were provided by Xu Cao (Johns Hopkins University School of Medicine). The mice were maintained in the animal facility at Indiana University School of Medicine. The animal experiment protocols were approved by the Institutional Animal Care and Use Committee at Indiana University School of Medicine in accordance with the National Institutes of Health Guide for the Care and Use of Laboratory Animals.

## METHOD DETAILS

### Animal experiments

Female C57BL/6 mice with CED and WT littermates at the age of 6 weeks (N = 6-8/group), 8 weeks (N = 7/group), and 12 weeks (N = 15/group) were used to determine EDL muscle-specific force (Figure 1A). Eight-week-old female mice were subcutaneously administered a vehicle (sterile PBS) or 5 mg/kg ZA (Novartis, Basel, Switzerland) three times a week for 4 weeks (Figure 1B). Seven-week-old female WT mice and mice with CED were given 5 mg/kg vehicle (WT, N = 15; CED, N = 16) or 5 mg/kg ZA (WT, N = 15; ZA, N = 16) three times a week for 10 weeks (Figure 2). Eleven-week-old female WT mice (N = 10/group) were given a vehicle, 2.5 mg/kg ZA once a week, or 5 mg/kg ZA three times a week for 8 weeks. Each animal experiment was conducted once. For all survival procedures, mice were anesthetized with a ketamine/xylazine cocktail. At the termination of the study, serum was collected from the mice via cardiac puncture, the mice were euthanized, and their bones and soft tissues were dissected and saved for processing.

### Radiography

ZA treatment response was monitored longitudinally via radiography using a digital x-ray imager (KUBTEC, Milford, CT). Images of mice were acquired in a prone position at 2.7× magnification at baseline prior to ZA treatment, 4 weeks after the initiation of treatment (midpoint), and at the termination of the study.

### DXA

Mouse BMD was measured longitudinally using a PIXI Mus II mouse densitometer (GE Lunar IIl; Faxitron, Tucson, AZ) as described previously (Waning et al., 2015). Blinded analyses of whole-body BMD and BMD of the tibia and femur metaphysis and lumbar vertebrae was performed. Because CED is characterized by diaphyseal bone dysplasia, entire tibiae and femora were analyzed to capture bone changes. Values were expressed as percentage change over baseline.

### Micro-CT

*Ex vivo* bone micro-CT imaging was performed to determine the quantity and quality of bone at the distal femur, proximal tibia, and L5 vertebra using a viva CT 40 (SCANCO Medical, Wayne, PA). High-resolution scans were acquired using a 17.5-μm^3^ isotropic voxel size, 55-kVp peak x ray tube potential, and 200-ms integration time and were subjected to Gaussian filtration.

### Trabecular bone analysis (tibia)

The trabecular bone microarchitecture was evaluated in the proximal metaphysis of the tibia 0.4 mm distal to the growth plate and extending distally for 1 mm. A threshold of 170 mg HA/cm^3^ was used to manually delineate bone from surrounding soft tissue. Parameters measured included trabecular bone volume fraction, trabecular number (mm^-1^), trabecular thickness (mm), trabecular separation (mm) and connectivity density, and structure model index.

### Trabecular bone analysis (L5 vertebra)

A 500-μm uniform core of trabecular bone, excluding the primary spongiosa, within the body of the L5 vertebra was evaluated for bone microarchitecture measurement. A threshold of 220 **mg** HA/cm^3^ for each bone slice was set to separate cortical and trabecular bone using an automated script.

### Cortical porosity analysis

The percent cortical porosity was determined in the tibia, where the region of interest was 15 mm from the midshaft to the ankle. Two-directional contouring was performed to exclude the medullary area. A threshold of 250 mg HA/cm^3^ was used to segment the bone from the medullary area and surrounding soft tissue. A separate analysis was performed to determine cortical parameters, such as cortical thickness (mm), medullary area (mm^2^), and cortical bone area (% BA/TA).

### Measurement of cross-sectional muscle areas

Proximal tibiae were scanned using a vivaCT40 to measure muscle cross-sectional areas. Both hindlimbs were scanned starting at the tibiofibular junction for 4-5 mm under general isoflurane based anesthesia. Scans were acquired using parameters of 45 kVp, 133 μA, and 620-ms integration time as a standard setting to optimize the contrast between muscle and fat tissue.

### Bone mechanical testing

Mechanical testing by compression was performed on L5 vertebrae and by three-point bending on femora from all groups of mice. Femora were tested in a three-point bending configuration with the anterior side facing down along 2 horizontal supports spaced 7 mm apart. The central loading point was displaced downward at 0.1 mm / s on the posterior surface of at the mid diaphysis. For all tests, load-displacement data were recorded at a sampling frequency of 100 Hz (Test Works 4.0). Curves were analyzed to measure whole-bone properties, primarily ultimate force, energy absorption to various points along the curve, stiffness, and displacement. Samples were tested in 37 μC of Hanks’ balanced salt solution in a three-point bending configuration with a custom-made rig for the ELF 3200 Series mechanical testing machine (BOSE Enduratec electromagnetic and mechanical testing devices;). L5 vertebral bodies were prepared with flat and parallel cranial and caudal ends by removing the intervertebral disc cartilage to expose the bone. Platen loading was performed at a rate of 3 mm / min.

### Bone histology and histomorphometry

At the time of euthanasia, mouse hindlimbs were dissected, fixed in 10% neutral-buffered formalin for 48 hours, and transferred to 70% ethanol. Tibiae were decalcified in 10% EDTA for 2 weeks, processed using an automated tissue processor (Excelsior; Thermoelectric, Waltham, MA). After embedding bones in paraffin, longitudinal midsagittal sections 3.5 μm thick were cut and stained with hematoxylin and eosin along with orange G and phloxine to visualize bone and with TRAP to visualize osteoclasts. The osteoclasts at the bone surface and total bone surface were quantified using BIOQUANT image analysis software (version 2018; Nashville, TN).

To determine the dynamic bone parameters, bones were double-labeled with calcein injectioned on days 3 and 10 prior to euthanasia to maintain a 7-day interval between the two injections. Undecalcified femora and lumbar vertebrae were dehydrated in graded alcohol (70-100%), cleared in xylene, and infiltrated and embedded with methyl methacrylate under a vacuum. Using an automated microtome (Microm HM 360; Thermo Fisher Scientific, Waltham, MA), 4.5-μm transverse sections were removed from the femoral diaphysis at the midsagittal and from the L1-L4 vertebrae and mounted unstained on standard microscope slides. Plastic embedded sections were used to measure the following dynamic parameters: bone formation rate/bone surface (mm^3^/mm^2^/d), mineral apposition rate (mm/d), and mineralizing surface (%). To differentiate mineralized and osteoid bone tissue, plastic-embedded femora and vertebrae were subjected to von Kossa staining and counterstaining with tetra chrome (MacNeal; Poly sciences, Warrington, PA) for improved contrast. Nonmineralized osteoid bone tissue was quantified by measuring the osteoid volume, surface, and width (μm). All analyses were performed using BIOQUANT image analysis software.

### Histological staining to assess morphological osteocyte changes

Osteocyte morphology was assessed by observing the integrity of the LCN in vehicle- and ZA-treated female WT mice and mice with CED, which was visualized using Ploton silver staining (Jauregui et al., 2016, Dole et al., 2017). Briefly, deparaffinized and hydrated sections of bones were incubated in two parts 50% silver nitrate and one part 1% formic acid in a 2% gelatin solution as described (Jauregui et al., 2016). Subsequently, silver nitrate-stained bone sections were washed in 5% sodium thiosulfate, dehydrated, cleared, and mounted. Images were obtained at 100× magnification using a Nikon Eclipse E800 bright-field microscope (Nikon Instruments, Melville, NY). The area spanned by the osteocyte lacunae and canaliculi was quantified as the lacuno-canalicular area. ImageJ was used for quantification and a threshold was applied to gray scale images for detecting silver-stained lacunae and canaliculi in the area of the bone in each image. Thus, the LCN area was calculated after normalizing to the total bone area in each image captured. LCN areas were evaluated in four animals per group, and four or five images of the cortical bone region in each animal were obtained.

### Forelimb grip strength

For each mouse, forelimb grip strength was measured by allowing the mouse to grab a wire mesh attached to a force transducer (Bioseb, Vitrolles, France) that records the peak force generated as the mouse is pulled by the tail horizontally away from the mesh. All mice were acquainted with the wire mesh a week prior to the actual assessment. Three repeated pulls were performed with a 5-second interval between each pull. The absolute grip strength was calculated and expressed in grams as an average of the peak force recorded from all three pulls.

### Whole muscle contractility

The EDL muscle was dissected from the hindlimb. Using a 4-0 silk suture, the tendons on the EDL muscle at the proximal and distal sites were tied, and the other end of the suture was tied with stainless steel hooks. Muscles were then mounted between a force transducer (model 407A; Aurora Scientific, Aurora, Ontario, Canada) and an adjustable hook. The muscles were immersed in a glass chamber containing O_2_/CO_2_ (95/5%) bubbled Tyrode’s solution (121 mM NaCl, 5 mM KCl, 1.8 mM CaCl_2_, 0.5 mM MgCl_2_, 0.4 mM NaH_2_PO_4_, 24 mM NaHCO_3_, 0.1 mM EDTA, 5.5 mM glucose). Each muscle was stimulated to contract using a supramaximal stimulus between two platinum electrodes. The force-frequency relationships were determined by triggering contraction using incremental stimulation frequencies (0.5-ms pulses at 1-200 Hz for 350 ms at supramaximal voltage). Between stimulations, the muscles were allowed to rest for 3 minutes. Specific force (kN/m^2^) was determined by normalizing the absolute force to the muscle cross sectional area with a muscle density constant of 1.056 kg/m^2^, thus correcting for differences in size. An endurance protocol to measure fatigue was run by inducing a tetanic response (70 Hz and 300 ms) and repeating for 50 cycles for the EDL muscle. Force and fatigue data were collected using the Dynamic Muscle Control/Data Acquisition and Dynamic Muscle Control Data Analysis software programs (Aurora Scientific).

### Immunoprecipitation and Western blotting

EDL muscles were lysed in a buffer containing consisting of 50 mM Tris-HCl (pH 7.4), 150 mM NaCl, 20 mM NaF, 1 mM Na_3_VO_4_, and protease inhibitors. Homogenized samples (150 μg) were incubated with an anti-RyR1 antibody in modified RIPA buffer (50 mM Tris-HCl, pH 7.4, 0.9% NaCl, 5 mM NaF, 1 mM Na_3_VO_4_, 1% Triton-X100, protease inhibitors) for 1 hour at 4°C. The immune complex was incubated with protein A-Sepharose beads (GE Healthcare) overnight at 4°C. The beads were washed twice using modified RIPA buffer. Samples were loaded on Bolt 4 to 12%, Bis-Tris mini gels and separated and transferred using a Bolt electrophoresis system and reagents (Invitrogen, Carlsbad, CA). Membranes were blocked using 5% milk in TBST to prevent nonspecific binding. After washing, membranes were incubated with anti-RyR (Yenzyme custom made; 1:2000) (Callaway et al., 1994) and anti-Cys-NO (cat# Ab94930; Abcam, Cambridge, UK; 1:2000) antibodies or an anti-calstabin antibody (cat# Ab2918; Abcam; 1:2500). EDL muscles were homogenized in NP-40 buffer to detect pSMAD3, SMAD3, Nox4 (cat# 109225; Abcam; 1:1000 each), GAPDH, and tubulin (Sigma, St. Louis, MO; 1:500 each). Immunoblots were developed and quantified using infrared-labeled secondary antibodies and an infrared imaging system (LI-COR Biosciences, Lincoln, NE). Levels of channel oxidation (DNP), S-nitrosylation (Cys-NO), and bound calstabin 1 (for RyR1 only) were normalized according to the total RyR1.

### Muscle histology, immunohistochemistry, and immunofluorescence

The RF muscle was dissected and fixed in 10% neutral buffered formalin for 24 hours, transferred to 70% ethanol, and processed using an automated tissue processor (Excelsior). After embedding in paraffin, longitudinal midsagittal sections, and cross-sections 8 μm thick were cut and stained with hematoxylin and eosin. Five to six fields (10×) per animal were analyzed to determine muscle fiber diameter and myofiber area using an automated muscle-specific script with BIOQUANT image analysis software.

Immunohistochemical analysis of phospho-SMAD2/3 was performed using paraffin embedded muscle sections. Briefly, tissue sections were deparaffinized and treated with 3% hydrogen peroxide to remove endogenous peroxidase. Sections were used for antigen retrieval with trypsin digestion (Invitrogen) followed by blocking of nonspecific binding sites using normal goat serum (10%) and an avidin/biotin blocking kit (Vector Laboratories, Burlingame, CA). Sections were incubated with phospho-SMAD2/3 (Cell Signaling Technology, Danvers, MA) at a 1:500 dilution overnight at 4°C. The next day, after washing with PBST, sections were incubated with a biotin-conjugated anti-rabbit antibody (EMD Millipore, Billerica, MA) and a streptavidin-peroxidase conjugate (Vector Laboratories) at a 1:1000 dilution for 1 hour at 37°C. Slideware stained using a 3,3-diaminobenzidine substrate kit (Vector Laboratories) and counterstained with hematoxylin. The percentage of phospho-SMAD2/3-positive nuclei per field was quantified. Five fields (20×) per animal were analyzed.

### Muscle fibrosis

Paraffin-embedded muscle sections were stained using Masson’s trichrome stain (HT-15 kit; Sigma) to differentiate collagen fibers, muscle fibers, fibrin, and erythrocytes. Briefly, deparaffinized and hydrated muscle sections were mordanted with preheated Bouin’s solution at room temperature overnight. The next day, slides were washed with running tap water to remove yellow color and stained using Weigert’s iron hematoxylin solution for 5 minutes followed by washing with water. Sections were placed in Biebrich scarlet-acid fuchsin, working phosphotungstic/phosphomolybdic acid solution and with aniline blue solution for 5 minutes in each solution with washes in between using deionized water. Slides were placed in an acetic acid solution followed by dehydration and mounted to visualize the blue-colored fibrosis in muscle sections.

For immunofluorescence assay, the RF muscle was dissected and placed on Tissue-Tek embedding medium (Sakura Finetek, USA) on a cork and flash-frozen in liquid nitrogen-cooled isopentane. Cryosections (10 μm) were cut using a cryostat and mounted on glass slides. Muscle sections were blocked using 10% normal goat serum in 1× PBS followed by overnight incubation in a moist chamber at 4°C with the following primary antibodies (DSHB, Iowa) diluted in a 10% normal goat serum/PBS solution: BA-F8 (1:25) or BA-D5 (1:100) for I, SC-71 (1:6) for IIA, BF-F3 (1:20) for IIB, and 05-206 (1:500) for laminin sections. Next day sections were washed with 10% normal goat serum and incubated with appropriate secondary antibodies for 1 hour at room temperature in a humidified chamber and with a secondary antibody cocktail (Life Technologies, Waltham, MA) of Alexa Fluor 546 (1:250), Alexa Fluor 594 (1:250), Alexa Fluor 488 (1:250), and Alexa Fluor anti-mouse 594 (1:500). After mounting, muscle images were acquired using a ZEN Pro image analysis system (Carl Zeiss Deutschland Microscopy GmbH, Oberkochen, Germany). Type I (blue), type IIa (red), type IIb (green), and type IIx (black) were counted, and the numbers of individual fibers were divided by the total number of fibers and expressed as percentages.

### ELISA

To determine serum TGFβ1 concentration, ELISA assay was performed using a commercially available Quantikine Mouse TGFβ1 ELISA kit (R&D Systems, Minneapolis, MN) according to the manufacturer’s guidelines. Serum samples were diluted as recommended by the manufacturer’s guidelines. Absorbance was read using a microplate reader at 450 nm with wavelength correction set at 540 nm.

### Mobility detection

Open field testing has been commonly used for behavioral testing. This test has been used by our group for mobility detection and activity monitoring in mice. In the present study, the open field test was performed in a chamber with a 48 cm × 48 cm square area and 25.5-cm-high walls. The square area was further divided into a 12 cm × 12 cm square, where the middle four squares were defined as the central zone and surrounding 12 squares were defined as the peripheral zone. A video tracking system was used to track the activity and locomotion of the mice. Each mouse was released in the center of the chamber and allowed to move freely in it. Movement traces of each mouse were recorded for 5 minutes using the video tracking system. The number of mouse entries to the center and peripheral quadrant of the chamber was automatically recorded and analyzed using ANY-maze software (Stoelting Co., Wood Dale, IL). Also, the elevated plus maze is commonly used to determine anxiety in mice, but we took advantage of this maze apparatus to monitor the movement of mice. This apparatus contains four arms: two without wall calls (open arms) and two with 15-cm-high wall calls (closed arms). This maze is 68 cm long and 5 cm wide and placed 55 cm off the floor at elevated position. The video tracking system was used to monitor mouse movements and time spent in the open and closed arms. Mice were released in the intersection of the four arms and allowed to move freely for 5 minutes. Their mobility and time were tracked using a video camera. The ratio of the number of entries in the open arm to that in the closed arm and the total time spent in the open and closed arms were analyzed. The apparatuses and software for both the open field and elevated plus maze tests were purchased from Stoelting Co.

### *In vitro* C2C12 cell experiments

C2C12 myoblasts (CRL-1772; ATCC, Manassas, VA) were cultured in Dulbecco’s modified Eagle’s medium (DMEM) containing 10% fetal bovine serum and a 1% penicillin-streptomycin solution and maintained at 37°C with 5% CO_2_ in a humidified chamber. Routine mycoplasma testing was performed using polymerase chain reaction. When C2C12 cells reached about 60% confluence, DMEM with 2% horse serum (Hy Clone Laboratories, Logan, UT) and 1% penicillin-streptomycin was added for differentiation. C2C12 cells were treated with ZA in the presence or absence of recombinant human TGFβ1 (R&D Systems).

### MTT assay

To determine ZA dose response in C2C12 cells, an MTT assay was performed using C2C12 myoblasts. Cells were seeded in 2% charcoal-stripped fetal bovine serum containing DMEM at a density of 250 cells per well in 96-well plates. Cells were treated with 5 ng/ml TGFβ1 in combination with ZA at six concentrations (0, 1.25, 2.5, 5, 50, and 100 μM) in DMEM with 2% stripped serum. Cells treated without TGFβ1 and ZA were used as controls. The media were changed every 24 hours. At the end point, MTT reagent (5 mg/ml; CalBioChem, San Diego, CA) was added to each well. After incubation for 4 hours at 37°C, cells were lysed using a PBS solution with sodium dodecyl sulfate (10%) and HCl (0.01 N), and the plate was incubated at 37°C for an additional 16 hours. The absorbance was measured at 570 nm using a plate reader.

### C2C12 fusion Index

For differentiation assays, differentiated C2C12 myotubes in six-well plates were treated with or without 5 ng/ml TGFβ1 and/or 0.625 and 1.25 μM ZA in 2% charcoal-stripped fetal bovine serum containing DMEM for 24 hours. C2C12 myotube cultures were fixed and permeabilized in an ice-cold solution of acetone: methanol (1:1) at -20°C for 20 minutes. Following fixation, the cells were rehydrated in PBS for 10 minutes at room temperature. Cells were then blocked with an 8% bovine serum albumin solution for 1 hour at room temperature, washed three times with PBS for 10 minutes, and incubated with a primary antibody against myosin heavy chain (R&D Systems) overnight at 4°C with gentle agitation. The following day, cells were washed with PBS (three times, 10 minutes each) and incubated with an Alexa Fluor 488-labeled anti mouse IgG (Life Technologies) secondary antibody for 1 hour at room temperature and protected from light, which was followed by three additional PBS washes. Nuclei were stained with DAPI for 1 minute followed by two PBS washes, and images were captured using an Axio Observer Z1 (Carl Zeiss Deutschland Microscopy GmbH). Myotube diameters, myotube/field numbers, average nucleus/myotube numbers, and the fusion index (ratio of the number of nuclei in a myotube to the total number of nuclei) were measured using ImageJ software (National Institutes of Health, Betehsda, MD).

### C2C12 RyR1 biochemistry

Differentiated C2C12 myotubes in a 10-cm dish were treated with or without 5 ng/ml TGFβ1 and/or 0.625 and 1.25 μM ZA in 2% charcoal-stripped containing DMEM for 24 and 48 hours. Differentiated C2C12 myotubes were lysed in NP-40 lysis buffer containing 50 mM Tris-HCl (pH 8.0), 150 mM NaCl, 1.0% NP-40, and protease inhibitors. Lysates were used for RyR1 precipitation similarly to that described for EDL muscle.

## Statistics

Statistical analyses were performed using Prism 7.0 software (GraphPad Prism, La Jolla, CA). Data presented as mean ± SEM. Unpaired Student’s *t* test, 2-tailed performed for comparisons involving two variables. A one-way analysis of variance (with a Tukey’s post-test) was performed for comparisons involving more than two conditions. *P* values of <0.05 were considered significant.

## SUPPLEMENTAL INFORMATION

**Figure S1. ZA dose response in muscle and bone of normal C57BL/6 mice**

Eight-week-old C57BL/6 mice were given a vehicle, 2.5 mg of ZA (once a week), or 5 mg of ZA (three times a week) for 4 weeks.

(A) Representative micro-CT reconstructions of trabecular bone at the proximal tibia.

(B) Quantification of trabecular bone volume at baseline (before treatment) and 4 weeks after treatment (N = 10/group).

(C) *Ex vivo* contractility of EDL muscle. EDL force was normalized with according to muscle size.

(D) Differences in hindlimb EDL, TA, soleus (SOL), and gastrocnemius (Gastroc) muscle weights in the three treatment groups.

Data are expressed as mean (± SEM) differences in muscle function determined using two-way ANOVA with Bonferroni’s multiple comparison testing. ***p < 0.001. Differences in all other parameters were determined using one-way ANOVA with Tukey’s multiple comparison test or an unpaired *t*-test. ***p < 0.001.

**Figure S2. Hindlimb muscle weights in WT mice and mice with CED treated with ZA for 10 weeks**

After 10 weeks of ZA-based treatment, hindlimb muscles were dissected and weighed. Differences in EDL, TA, soleus (SOL), and gastrocnemius weights are presented.

Data are expressed as mean (± SEM) differences in muscle function. Differences in all other parameters were determined using one-way ANOVA with Tukey’s multiple comparison test. *p < 0.05 and **p < 0.01. Differences between vehicle-treated WT mice and mice with CED and between vehicle- and ZA-treated mice with CED are presented.

**Figure S3. Skeletal muscle fibrosis**

RF muscle sections were stained using Masson’s trichrome stain. Muscle sections were scanned at 5× magnifications using BIOQUANT image analysis software (N = 6/group). Fibrotic tissues were stained blue, and muscle fibers were stained red as indicated by the arrows. Percent fibrosis was determined by measuring the percent fibrosis area by the total muscle area. Differences in fibrosis according to treatment and mouse type are presented in the graph.

**Figure S4. Skeletal muscle fiber type and mobility detection in mice**

(A) Fiber typing using TA muscle. Frozen TA muscle sections processed using isopentane were used for immunofluorescence with specific antibodies against type I, type IIa, type IIb, and myosin heavy chain fibers. **{Slow**-twitch fibers are shown in blue, intermediate fibers (between type I and IIa) are shown in red, fast-twitch (IIb) fibers are shown in green, and intermediate fibers (between IIa and IIb) are shown in black. All fibers were separated using laminin immunostaining. On each bar in the graph, a = type I versus types IIa, IIb, and IIx; b = type IIa versus type IIb; and c = type IIb versus type IIx. The colors of the letters correspond to the colors of the groups. Six TA muscle samples per group were used for immunofluorescent staining and analysis. *<0.05.

(B) Mobility detection using open field and elevated plus maze testing. Mice were placed in the center of an open field apparatus, and their activity was monitored using a video camera attached to the apparatus. Also, mice were released in the middle of the cross point of the elevated plus maze, and their activity was monitored using a video camera. In both apparatuses, the activity and locomotion of each mouse was monitored for 5 minutes. ANY-maze software was used to determine the number of mouse entries and time spent in particular area of the apparatuses.

**Figure S5. Effects of combination treatment with TGFβ and ZA on proliferation of, differentiation of, and RyR1 oxidation in undifferentiated and differentiated C2C12 cells**

(A) In the presence and absence of TGFβ, undifferentiated C2C12 myoblasts were treated with five concentrations of ZA to determine the optimum treatment dose. Also, C2C12 cells were treated with ZA for 72 hours to determine the cell growth rate using an MTT assay. Differences in the ZA treatment groups of mice are compared with those in a no treatment group.

(B) The differentiation potential of C2C12 myotubes in response to treatment with TGFβ and/or ZA calculated using the fusion index, number of myotubes, average number of nuclei per myotube, and myotube diameter. Comparisons of the TGFβ-positive and -negative groups and TGFβ-positive no ZA, 0.625 mM ZA, and 1.25 mM ZA groups are presented.

(C) Differentiated C2C12 myotubes were treated in the presence or absence of TGFβ and two concentrations of ZA for 24 hours followed by immunofluorescent staining using an anti-myosin heavy chain antibody and DAPI. Myotubes are shown in green, and sarcomeres are shown in blue.

(D) Differentiated C2C12 myotubes treated with or without TGFβ in the presence and absence of ZA were homogenized. An immunoblot of phosphorylation of SMAD3, co-immunoprecipitation of RyR1 oxidation and nitrosylation, Nox4, and calstabin 1 binding is shown.

Data are expressed as mean (± SEM) differences in MTT assays results determined using two way ANOVA with Bonferroni’s multiple comparisons testing. *p < 0.05 and ***p < 0.001.

Differences in all other parameters were determined using one-way ANOVA with Tukey’s multiple comparison test or an unpaired *t*-test.

