## Supplemental figure for "Zoledronic acid improves bone quality and muscle function in a high bone turnover state"

**Supplementary Figure 1**

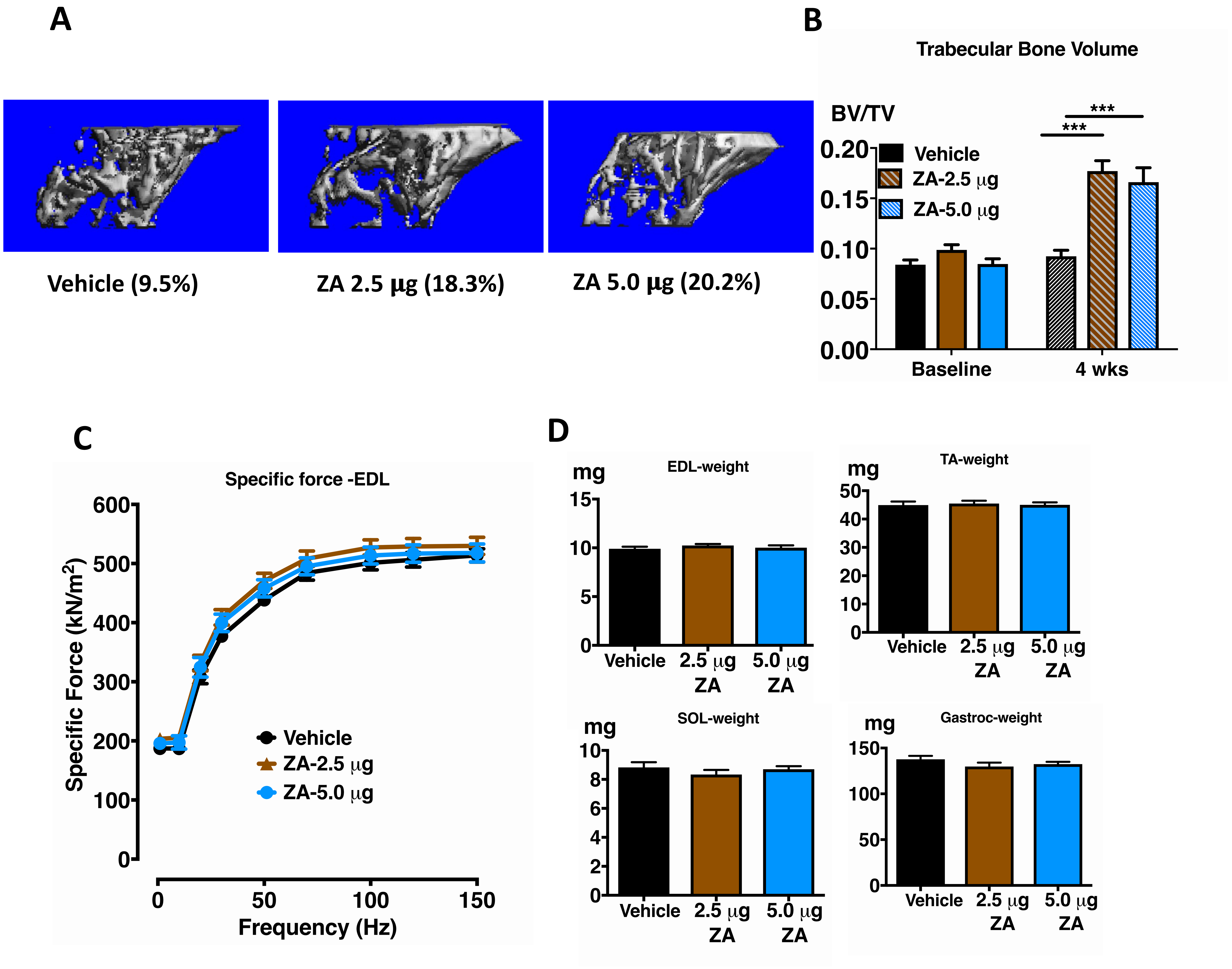

**Supplementary Figure 2**

### Muscle weights

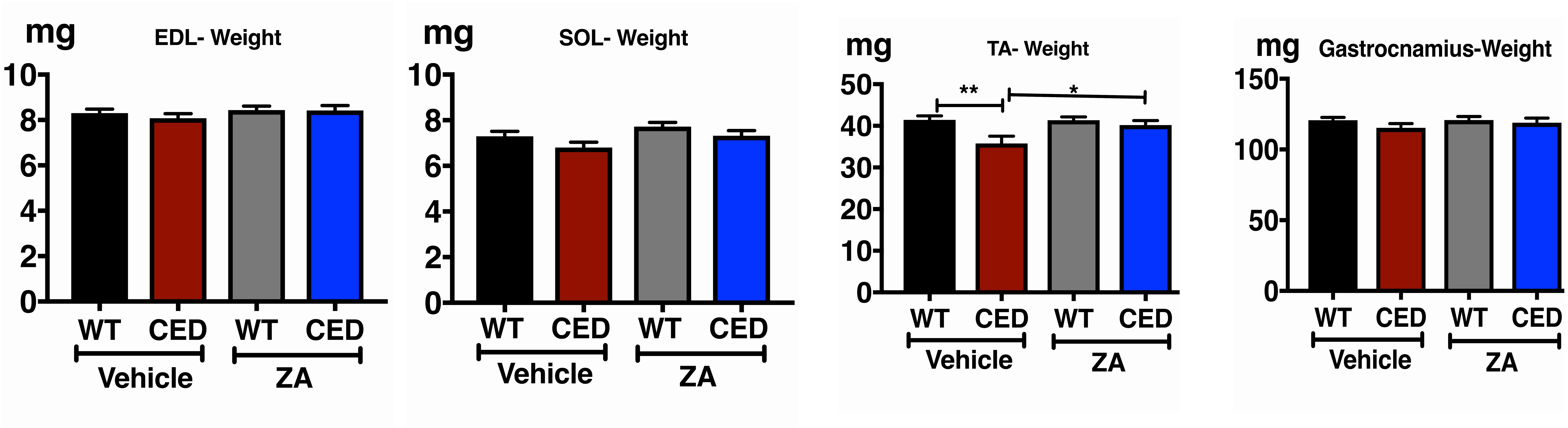

Supplementary Figure 3

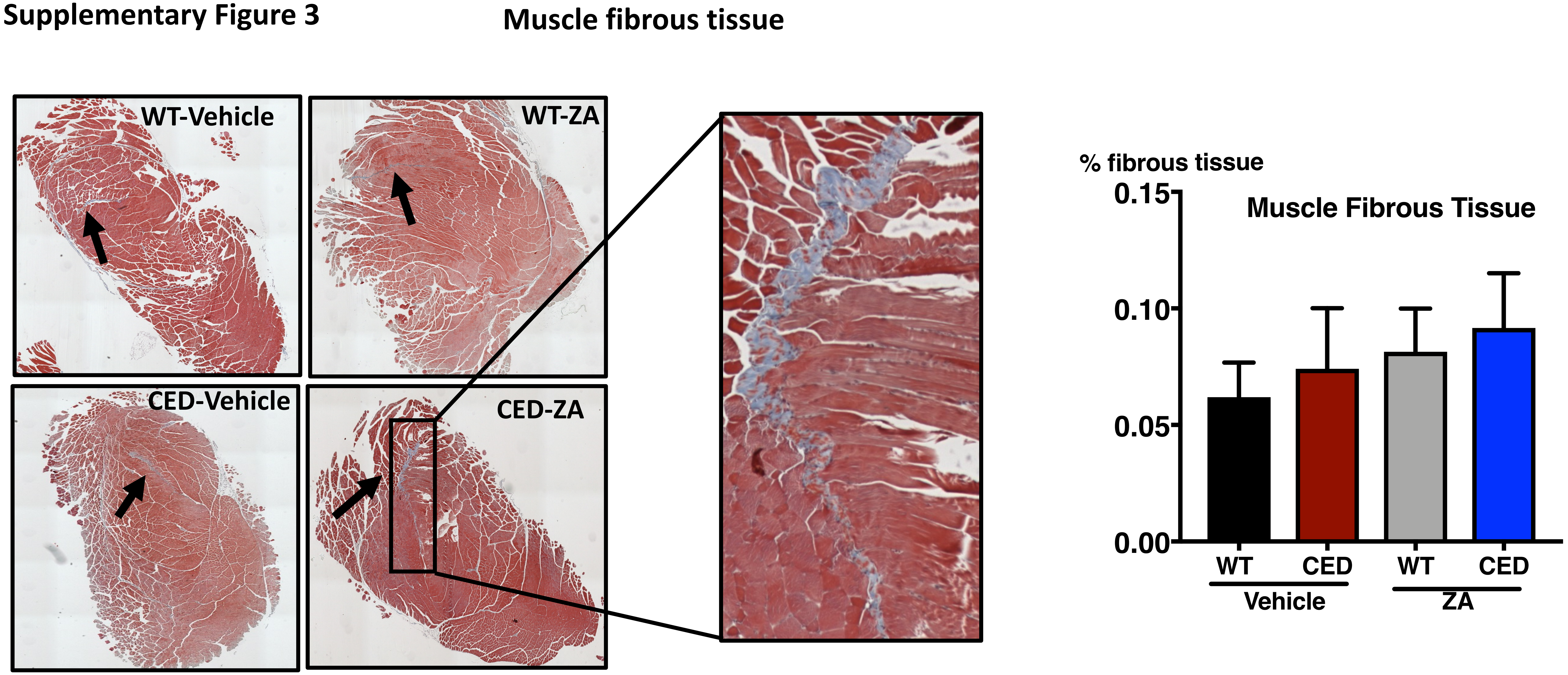

Supplementary Figure 4

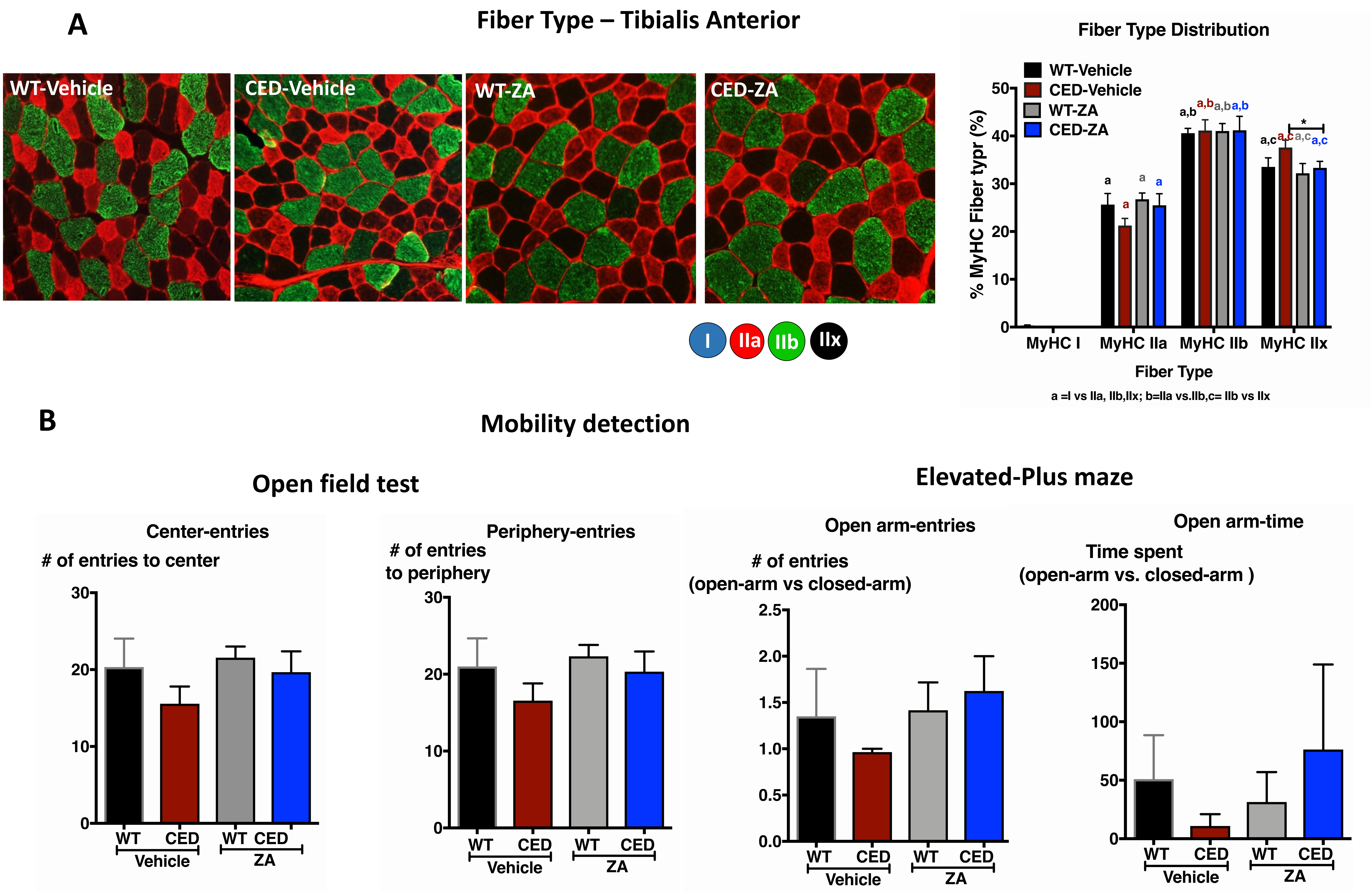

### Supplementary Figure 5

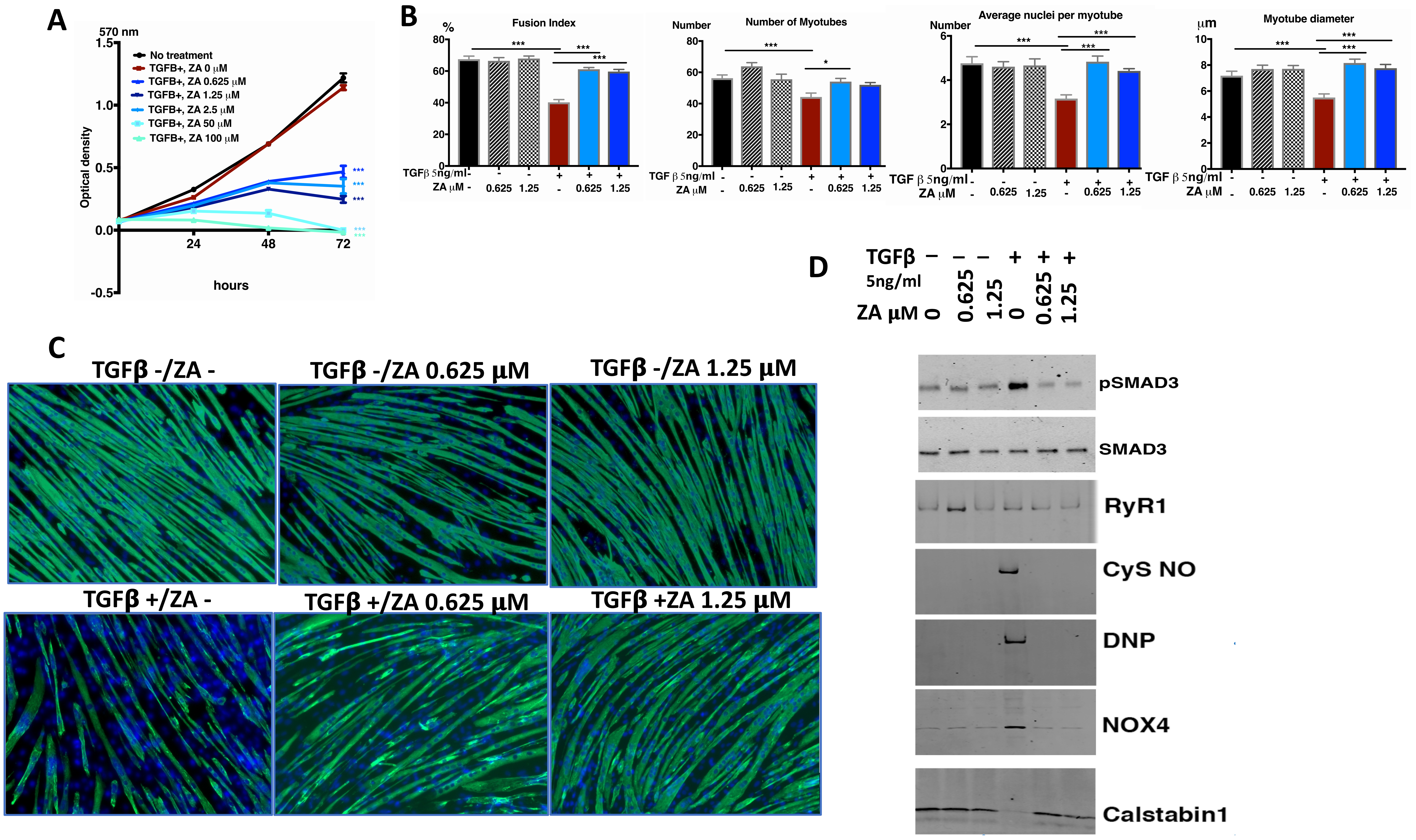
